# Human neonatal CITE-seq atlas identifies an immune transition at 32 weeks’ gestation from CD15⁺ myeloid-dominated to interferon-primed immunity

**DOI:** 10.64898/2026.04.01.715643

**Authors:** Paula Rothämel, Alessandro Mattia, Maria Isabel Corey, Barbara Puzek, Jana Wiesel, Pauline Michael-Kuschel, Christoph Klein, Markus Sperandio, Philipp Henneke, Claudia Nussbaum, Sarah Kim-Hellmuth

## Abstract

The human neonatal immune system is developmentally specialized to balance the unique requirements of perinatal transition. Disruption of this finely tuned balance, as in preterm birth, may have profound consequences for immunity and overall health. However, the impact of prematurity on immune composition and functional responsiveness across gestational ages (GA) remains incompletely understood. Single-cell profiling has advanced our understanding of neonatal immunity, yet most studies were limited to unimodal readouts, narrow GA windows, or baseline function. Here, we present a comprehensive human neonatal CITE-seq atlas (82 samples from 25 neonates and 10 adults as controls) at the first days of life covering a wide GA range and integrating baseline and stimulated conditions. Most notably, we identify a GA-dependent immune transition point centered around 32 weeks of GA, which discriminates extremely and very preterm neonates (GA <32wks) from those of higher GA (≥32wks). In particular, early-life immunity in extremely and very preterm infants showed CD15^+^ granulocytic myeloid derived suppressor cell-like predominance, whereas more mature neonates exhibited interferon-primed transcriptional profiles. This resulted in divergent myeloid-to-lymphocyte signaling networks and qualitatively distinct NK- and T-cell bystander responses upon activation. Together, these findings show that intrauterine development imprints GA-specific immune programs. By defining a developmental transition around a GA of 32 weeks that regulates baseline and induced responses of neonatal immune cells, our atlas provides a framework for understanding the vulnerability of preterm infants and thus may pave the way for developing GA-adapted immunomodulatory strategies.

**GRAPHICAL ABSTRACT:** 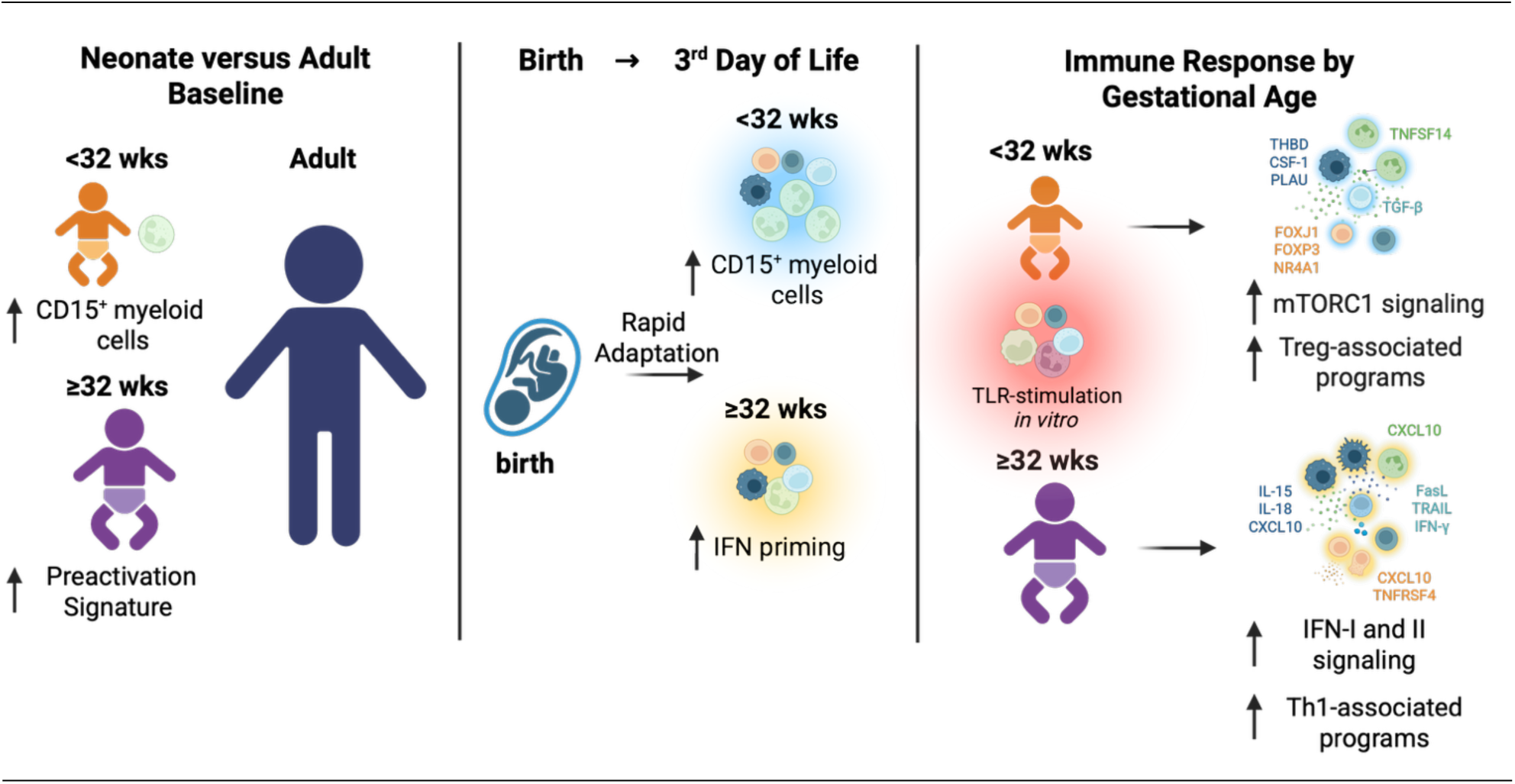

## INTRODUCTION

The neonatal immune system is increasingly recognized to be in a developmentally adapted state optimized to meet the unique challenges of perinatal transition. At birth, it must rapidly mount an effective response to microbial exposure while simultaneously preventing excessive inflammation. This balance is achieved through regulatory mechanisms that integrate immune tolerance and tissue protection with rapid responsiveness in a context-dependent manner, defining early-life immunity as distinct rather than deficient or uniformly attenuated.^1–3^

Preterm birth profoundly challenges this finely tuned immune program. Prematurity is associated with heightened susceptibility to bacterial and viral infections, reflecting an altered immune function.^4–5^ Beyond acute vulnerability, prematurity may induce long-lasting immune dysregulation, with evidence linking early-life immune perturbations to increased risk of inflammatory disease later in life.^6–8^ Longitudinal studies have demonstrated that term infants follow a relatively stereotypic postnatal immune trajectory,^8^ whereas extremely preterm infants diverge from this during the first months of life.^9^ These observations suggest that intrauterine development shapes postnatal immune programs; however, the underlying mechanisms remain incompletely understood. In particular, it remains unclear whether prematurity reflects a delay in maturation or the persistence of a quantitively and qualitatively distinct fetal immune program that leads to an altered postnatal immune phenotype.

A major knowledge gap lies in defining the specific cellular players and molecular programs that govern this developmental transition. In healthy term infants, immune adaptation involves a shift from a tolerogenic fetal state to a more postnatal configuration with heightened responsiveness. Central to this process are neonatal myeloid populations, including low-density neutrophils (LDNs) and their subset granulocytic myeloid-derived suppressor cells (gMDSCs), which are enriched during pregnancy and in cord blood and dampen excessive inflammation via mechanisms such as arginase-1 activity. These anti-inflammatory immune cell populations decline quickly after birth in the blood, suggesting that they may represent an important element in immune development.^10–13^ In parallel, term infants undergo an early transient hyporesponsive phase, followed by the establishment of a tonic interferon-signature during the first week of life that is associated with enhanced antiviral immunity throughout infancy.^14–16^ It remains unclear whether this reflects a gestationally timed developmental transition in human immune ontogeny.

Adaptive immunity is similarly developmentally structured. Neonatal T cells preferentially differentiate into regulatory T cells (Tregs) upon T cell receptor (TCR) engagement^17–18^ and exhibit a Th2-biased phenotype compared to adults.^19^ At the same time, they retain a pronounced capacity for rapid, TCR-independent bystander activation in response to innate cytokines, prioritizing early host defense over memory formation.^3, 20, 21^ How GA influences the balance between regulatory and effector programs in this context, and whether these pathways are differentially engaged in preterm infants, remains unknown.

Dissecting these complex, GA-dependent programs requires approaches capable of resolving cellular heterogeneity, activation states, and signaling programs, as well as their dynamic reorganization in response to inflammatory challenges at high resolution. Although single-cell profiling has transformed our understanding of neonatal immunity, most existing datasets rely on unimodal readouts or baseline measurements alone, limiting the detection of rare, transitional, or functionally defined cell states and their intercellular signaling programs in the rapidly evolving neonatal immune system.

To address these gaps, we systematically characterized circulating immune cells from 25 neonates spanning a broad GA spectrum (24-42 weeks) and 10 adults using a multimodal Cellular Indexing of Transcriptomes and Epitopes by Sequencing (CITE-seq) approach. We examined baseline immune organization during the first days of life and assessed GA-dependent responses to stimulation of pattern recognition receptors *in vitro*. Specifically, we asked: (i) how do immune cells differ between neonatal and adult samples under baseline conditions; (ii) how does the perinatal immune system evolve within the first three days of life; and (iii) how does GA shape neonatal immune responses to innate stimulation *in vitro*. Our analyses reveal that GA structures early-life immune organization, with a marked developmental transition around 32 weeks of gestation. Preterm neonates born before this point exhibit a CD15^+^ gMDSC-like dominated immune phenotype, whereas increasing GA is associated with the emergence of interferon-primed transcriptional states and distinct patterns of myeloid-lymphoid communication and NK- and T-cell bystander responsiveness.

To our knowledge, this study provides the first systematic CITE-seq comparison of human neonatal immune cell states across GA in the immediate postnatal period, integrating both baseline and innate stimulation conditions. By defining a GA-dependent developmental transition that stratifies immune architecture and functional responsiveness, our atlas establishes a framework for understanding immune vulnerability in extremely and very preterm infants.

## RESULTS

### Neonatal CITE-seq atlas reveals separation by condition and gestational age at 32+0 weeks

We enrolled 25 neonates at the LMU University Hospital neonatal intensive care unit from different gestational age categories, classified according to the World Health Organization (WHO) definition of preterm birth, including extremely preterm (<28+0 weeks of gestation, n=5), very preterm (28+0 to 31+6 weeks, n=2), late preterm (32+0 to 36+6 weeks, n=5) and full-term infants (≥37+0 weeks, n=13). Blood was collected at birth (n=13) and/or on the 3^rd^ day of life (DOL3) (n=12). The preterm infants had complications that were typical for their degree of prematurity, including one neonate with clinical evident chorioamnionitis; however, none of the included infants fulfilled the criteria for early-onset sepsis (Tables S1-S7). Neonatal peripheral blood mononuclear cells (PBMCs) and cord blood mononuclear cells (CBMCs) were either left untreated (baseline) or treated for 2 hours with Lipopolysaccharide (LPS, 200 ng/mL) or the TLR7/8 ligand Resiquimod (R848, 500 ng/mL), respectively. In addition, baseline PBMCs from 10 healthy adults were used as controls. In total, 195.000 cells of 82 samples from 35 individuals were profiled using a 36-antibody CITE-seq panel (Tables S8-S9) (Figure 1A). Principal component analysis (PCA) of normalized RNA expression, aggregated by donor and condition, demonstrated a clear separation between neonatal and adult samples (R^2^=0.53, p < 0.001) and, notably, a stratification by gestational age (GA) within neonates, separating extremely and very preterm (<32+0-wk) from late preterm and full-term (≥32+0-wk) neonatal samples (Figure 1B). This stratification was supported by PERMANOVA analysis (R^2^=0.45, p < 0.001), indicating a strong association between GA and transcriptomic variation, although differences within-group variability also contributed to this pattern. This separation was further accentuated following TLR stimulation, indicating that the induced transcriptional response of inflammatory genes was dependent on the GA. Consistent with this, PCA restricted to baseline samples confirmed a robust segregation between neonatal and adult samples (Figure S1A). Accordingly, we first characterized global neonatal vs. adult differences and subsequently examined GA-dependent (<32+0-wk vs. ≥32+0-wk) heterogeneity within the neonatal cohort. Multimodal integration of scRNA-seq and antibody-derived tag (ADT) data at baseline yielded 17 cell type clusters (Figures 1C, S1B, and S1C).

**Figure 1.**
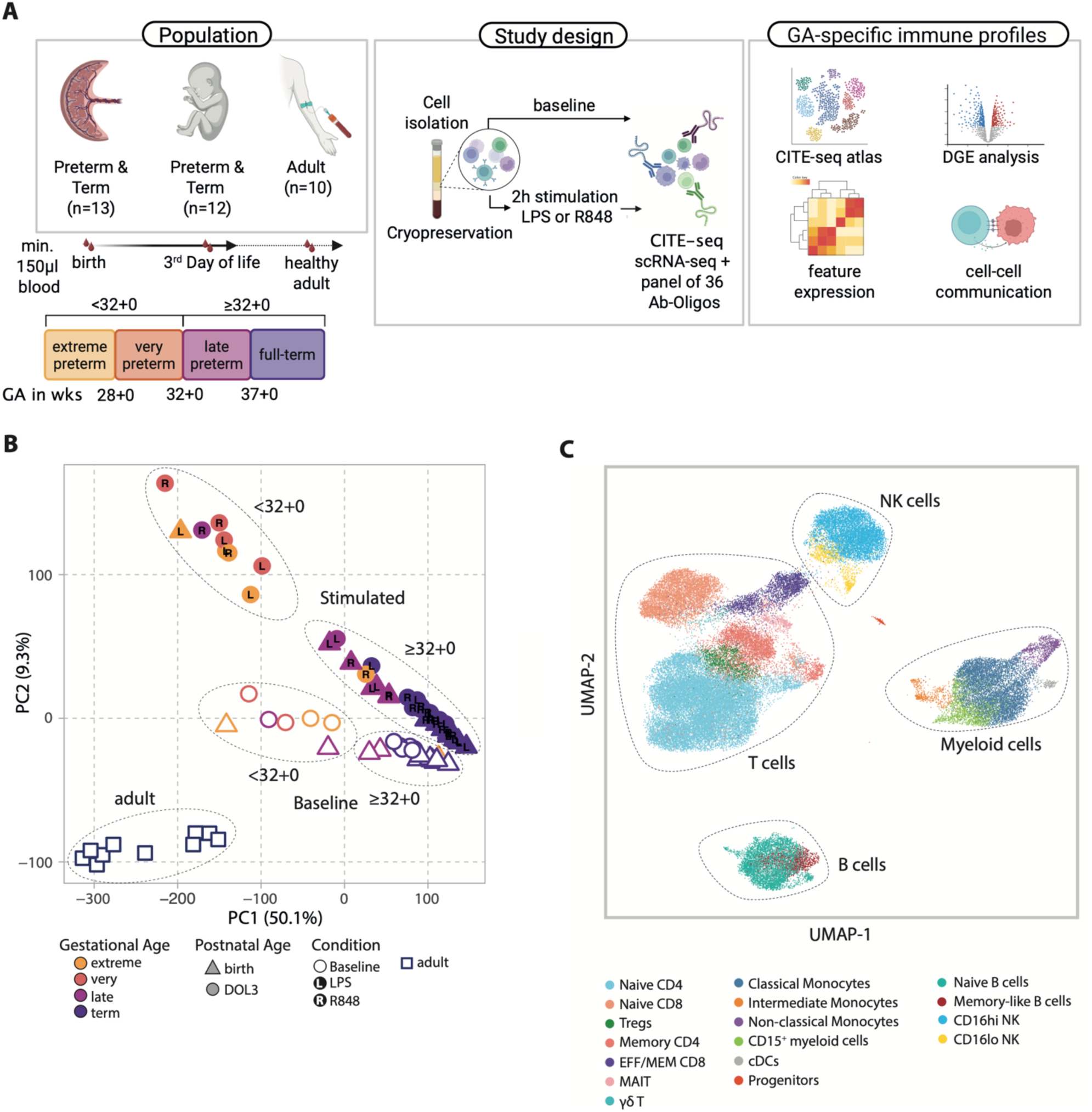
Neonatal CITE-seq atlas reveals separation by gestational age and condition. (A) Schematic of the experimental design showing patient groups and processing steps (created with BioRender). CITE-seq data generated from preterm and term circulating immune cell samples, collected right after birth or on the 3^rd^ day of life, or from healthy adult donors (82 samples from 25 neonates and 10 adults), either left untreated or following stimulation with TLR agonists (LPS or R848). (B) PCA of normalized RNA expression, aggregated by donor and condition. (C) UMAP plot of peripheral immune cell subsets identified within unstimulated neonatal and adult samples. wks, weeks; PBMC, peripheral blood mononuclear cells; LPS, Lipopolysaccharide; extreme, extremely preterm (<28+0 weeks of gestation); very, very preterm (28+0 to 31+6 weeks); late, late preterm (32+0 to 36+6 weeks); term, full-term (≥37+0 weeks).

### Neonatal vs. adult innate immune profiling at baseline reveals enrichment of ARG1_high_ CD15^+^ myeloid cells in <32+0-wk preterm infants

To systematically characterize immune profile differences between neonatal (n=25) and adult (n=10) individuals under baseline conditions (Figure S1A), we compared cell type clusters within major immune cell types, including myeloid, NK, T, and B cell subsets. Given the particular developmental adaptations assigned to innate immunity in early life,^14,22^ we first focused our analysis on the neonatal myeloid and NK cell compartment to delineate transcriptional and compositional differences compared to adults.

Clustering of the myeloid cell compartment identified five myeloid populations, including classical, intermediate, and non-classical monocytes, as well as conventional dendritic cells (cDCs) (Figure 2A). Notably, incorporating ADT surface protein expression, enabled the identification of CD15^+^ myeloid cells that separated into two distinct subclusters (Figures S2A and S2B). As conventional neutrophils localize to the polymorphonuclear (PMN) fraction during density gradient centrifugation, CD15^+^ cells identified within PBMCs likely represent low-density neutrophils (LDNs). One cluster was enriched for pro-inflammatory genes (e.g. *CXCL2*, *CXCL8,* and *IL1B*). The other cluster expressed *ARG1*, *ALPL*, and *CCL4*, markers previously linked to granulocytic myeloid-derived suppressor cells (gMDSCs; Figure S2B) and cells of this cluster are referred to hereafter as ARG1_high_ CD15^+^ myeloid cells. CD15^+^ myeloid cells were proportionally enriched in neonates; however, there were distinct GA-dependent differences between the subclusters. Extremely and very preterm individuals (<32+0-wk) displayed mostly CD15^+^ myeloid cells with a gMDSC-like transcriptional signature. In contrast, more mature neonates (≥32+0-wk) were characterized by CD15^+^ myeloid cells with the pro-inflammatory phenotype. Bayesian compositional analysis^23^ showed a significant increase of ARG1_high_ CD15^+^ myeloid cells and decreased classical monocytes in <32+0-wk compared to ≥32+0-wk and adults, respectively (Figures 2B, 2C, S2C, and S2D). Flow cytometry analyses of neonatal (n=22) and adult (n=7) PBMC samples obtained between postnatal days 2-4 confirmed a higher frequency of CD15^+^ myeloid cells with granulocyte-like scatter characteristics among viable cells in neonates, most likely corresponding to LDNs, with significantly higher frequencies observed in <32+0-wk compared to ≥32+0-wk neonates and adults (Figures 2D, 2E, and S2E, Tables S10-S11). In addition, CD15⁺ myeloid cells showed marked heterogeneity in CD16 surface expression (Figure 2D). CD16 expression levels have previously been linked to functional specialization among LDN populations, with only CD16_high_ LDNs demonstrating immunosuppressive properties characteristic of gMDSCs.^24^ ARG1_high_ CD15^+^ myeloid cells in our CITE-seq dataset were also CD16_high_, consistent with a gMDSC-like phenotype (Figure S2A).

**Figure 2.**
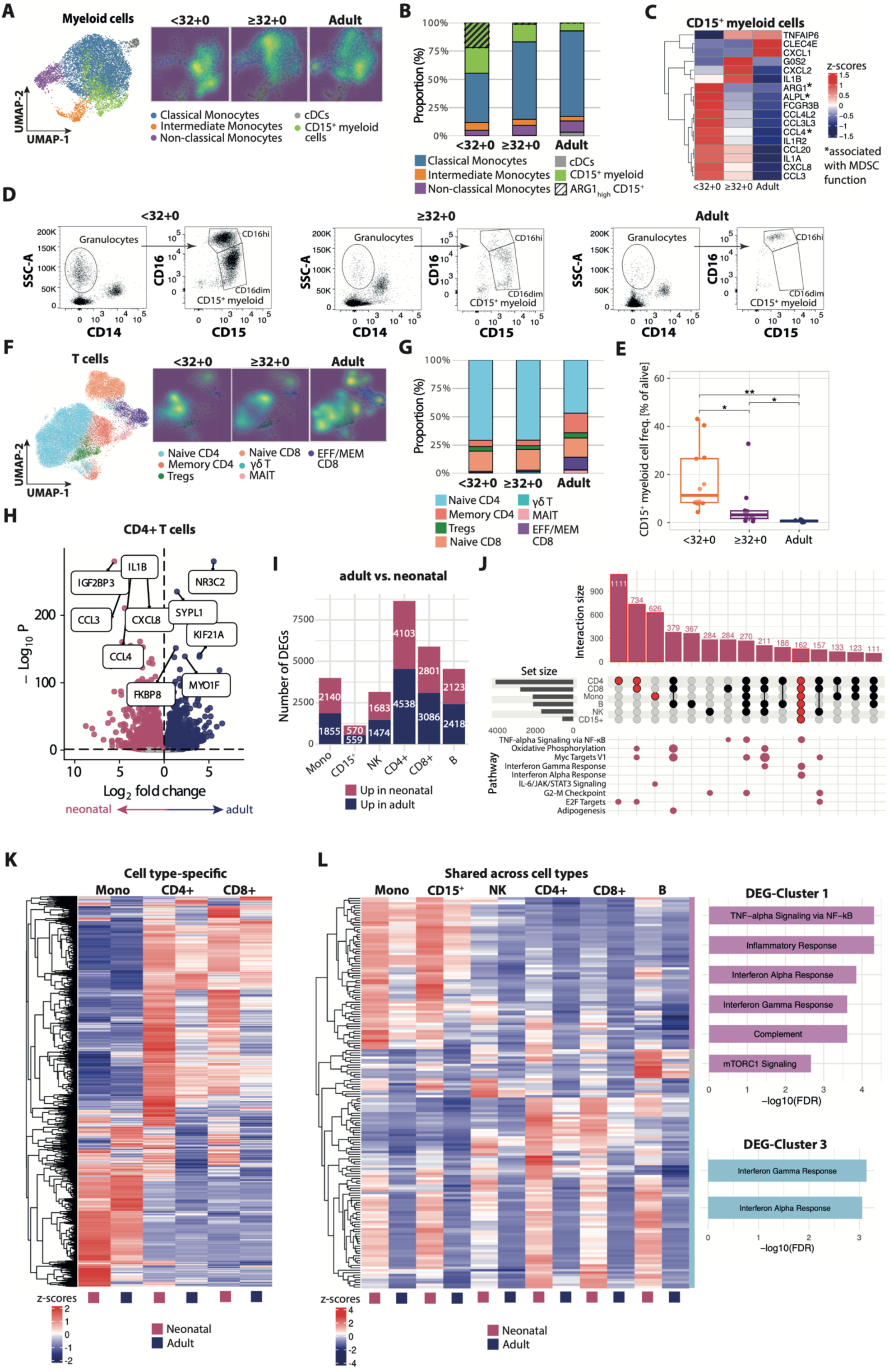
Increased CD15^+^ myeloid cell proportion and signs of basal preactivation in neonates compared to adults. (A) Left: UMAP plot of myeloid cells at baseline, derived from neonatal birth and DOL3, as well as adult samples. Right: Normalized density UMAPs of myeloid cells split by age group (neonates born <32+0 weeks of gestation, neonates born ≥32+0 weeks of gestation, and adults). (B) Stacked bar graph representing relative proportion of cells per cell type and group among myeloid cells. (C) Heatmap of selected marker genes for CD15^+^ myeloid cells, grouped by age group. (D) Representative flow cytometry plots from a very preterm (<32+0-wk), term (≥32+0-wk), and adult PBMC sample illustrating the distribution of CD15⁺ myeloid subsets. (E) Frequency of CD15⁺ granulocytic myeloid cells among alive cells across age groups as assessed by flow cytometry (<32-wk, n=12; ≥32+0-wk, n=10; adults, n=7; pairwise Wilcoxon rank-sum tests; p_Bonf_: * < 0.05, ** < 0.01). (F) UMAP and normalized density plots of T cells at baseline across age groups, as in (A) for myeloid cells. (G) Relative proportions of T cell subsets across age groups, as in (B) for myeloid cells. (H) Volcano plot showing differential gene expression between adult and neonatal CD4+ T cells at baseline. Pink, upregulated in neonatal; blue, upregulated in adult (adj. p < 0.05). Top DEGs (|log2FC| > 1) were labeled based on ranked adjusted p values. (I) Stacked bar plot of differentially expressed gene counts by cell type (adj. p < 0.05). (J) UpSet plot showing intersections of genes upregulated in neonatal samples (adj. p < 0.05) across six immune cell types. Bars indicate the number of genes shared between cell types. (K) Hierarchically clustered heatmap showing cell type-specific expression patterns of genes identified in (J). (L) Left: Hierarchically clustered heatmap of 162 genes consistently upregulated in neonatal vs. adult across different cell types as identified in (J) (cutree=3). Right: Enrichment analysis of DEG clusters using enrichR (MSigDB); top enriched terms ranked by FDR; FDR < 0.05.

In the NK cell compartment, we observed pronounced GA-dependent patterns across NK subclusters, most prominently within CD16hi NK cells (Figure S2F). We further stratified CD16hi NK cells and identified one subset (Cluster I) characterized by high expression of *CD8A*, *CX3CR1*, and *TXNIP* (Figure S2G). Notably, CD8 was also detected at the ADT level, highlighting another unique advantage of CITE-seq enabling cluster identification based on both transcriptional programs and surface marker expression (Figure S2H). This CD8_high_ cluster was proportionally enriched among CD16hi NK cells from <32+0-wk (Figure S2I). This finding was independently confirmed by flow cytometry in a separate group of 31 neonates (PBMCs collected at a median postnatal age of 3 days [range, 1-37]) and 9 adults, showing a higher frequency of PBMC-derived CD8+ NK cells in <32+0-wk compared to ≥32+0-wk (Figures S2J and S2K). The co-expression of *CX3CR1* and *TXNIP* is notable (Figure S2G), as both genes have been associated with NK cell states showing reduced IFN-γ production.^25,26^ Moreover, CD8+ NK cells themselves have previously been described as exhibiting exhausted or regulatory features,^27,28^ supporting the interpretation that this cluster reflects a distinct NK cell state with features indicative of altered activation or effector capacity.

### Neonatal adaptive immune cells show signs of basal preactivation

Beyond innate immune cell subsets, neonatal T cells are increasingly recognized as uniquely adapted to the immunological demands of early life.^21,29^ To investigate T cell phenotypic and transcriptional dynamics, we next analyzed the blood circulating T cell compartment in detail. Compared to adults, most neonatal T cells displayed a naïve phenotype (Figures 2F and 2G), a pattern also observed among neonatal B cells (Figures S2L and S2M). While overall proportions did not differ markedly between extremely and very preterm neonates <32+0-wk and more mature infants ≥32+0-wk, subcluster analysis revealed distinct age-dependent patterns, particularly within CD4+ T cells (Figure 2F, light blue cluster). Subsequent differential expression analysis showed upregulation of inflammatory and chemokine genes in neonatal CD4+ T cells relative to adults, including *CCL4, CCL3, IL1B,* and *CXCL8* (Figure 2H). Thus, neonatal T cells exhibited a transcriptomic profile indicative of a relatively preactivated baseline state.

### Pathways related to IFN-I and II and TNF-α signaling are enriched at baseline in neonatal immune cell subsets

Analysis of differentially expressed genes (DEGs) in neonatal CD4+ T cells revealed extensive transcriptional changes compared to adults, prompting a systematic examination of DEGs across all major immune cell types, including monocytes, CD15^+^ myeloid cells, NK cells, T cells, and B cells. Overall, the number of DEGs varied by cell type: CD4+ T cells exhibited the highest number (4103 up- and 4538 downregulated genes, respectively), whereas CD15⁺ myeloid cells showed the lowest number (570 up- and 559 downregulated genes, respectively), consistent with their intrinsically lower transcriptional activity, as expected for a neutrophil-like population (Figure 2I).

Among genes upregulated in neonates, the majority displayed cell type-specific expression patterns (Figures 2J and 2K, Table S16). For example, genes selectively upregulated in neonatal T cells were enriched for E2F target gene signatures, oxidative phosphorylation, and MYC target pathways (Figure 2J). In contrast, genes specifically upregulated in neonatal monocytes were enriched for IL-6/JAK/STAT3 signaling (Figure 2J), highlighting lineage-specific transcriptional programs within the neonatal immune system.

In addition to these cell type-specific signatures, a subset of genes was commonly upregulated across all major neonatal immune cell types (Figure 2J). These shared genes were enriched for pathways related to TNF-α signaling via NF-κB, inflammatory responses, and interferon-α and -γ signaling (Figure 2L). Although individual DEG clusters showed stronger expression in particular neonatal cell types - such as pro-inflammatory TNF-α/NF-κB-associated genes in myeloid cells (DEG-Cluster 1) and interferon-responsive genes in CD4+ T cells (DEG-Cluster 3) - their presence across multiple cell types may reflect broader immune activation patterns in neonatal immunity.

Overall, compared to adults, neonatal samples exhibited enrichment of subsets with potentially regulatory functions - including ARG1_high_ CD15^+^ gMDSC-like cells and CD8_high_ NK cells, particularly in extremely and very preterm infants <32+0-wk. In addition, they exhibited widespread signs of preactivation across most cell types. This raises important questions about whether this activation is already established at birth and how it is modulated by prematurity.

### Immune cells from ≥32+0-wk neonates exhibit interferon priming on DOL3, whereas <32+0-wk show persistence of CD15^+^ myeloid dominance

To determine whether basal immune activation is present at birth or evolves during the early postnatal period in a GA-dependent manner, we compared immune profiles at birth and at DOL3, examining both cell compositional and transcriptional differences across all major immune cell types. Within the myeloid compartment, the proportion of CD15^+^ myeloid cells tended to increase from birth to DOL3 in extremely and very preterm neonates <32+0-wk, whereas in more mature neonates ≥32+0-wk, it declined toward a more adult-like cell composition. Notably, ARG1_high_ CD15⁺ gMDSC-like cells remained abundant in <32+0-wk (Figure 3A). In contrast, the relative proportion of NK, T, and B cell subsets showed only modest GA-related variation over the same interval (Figures 3B, S3A, and S3B).

**Figure 3.**
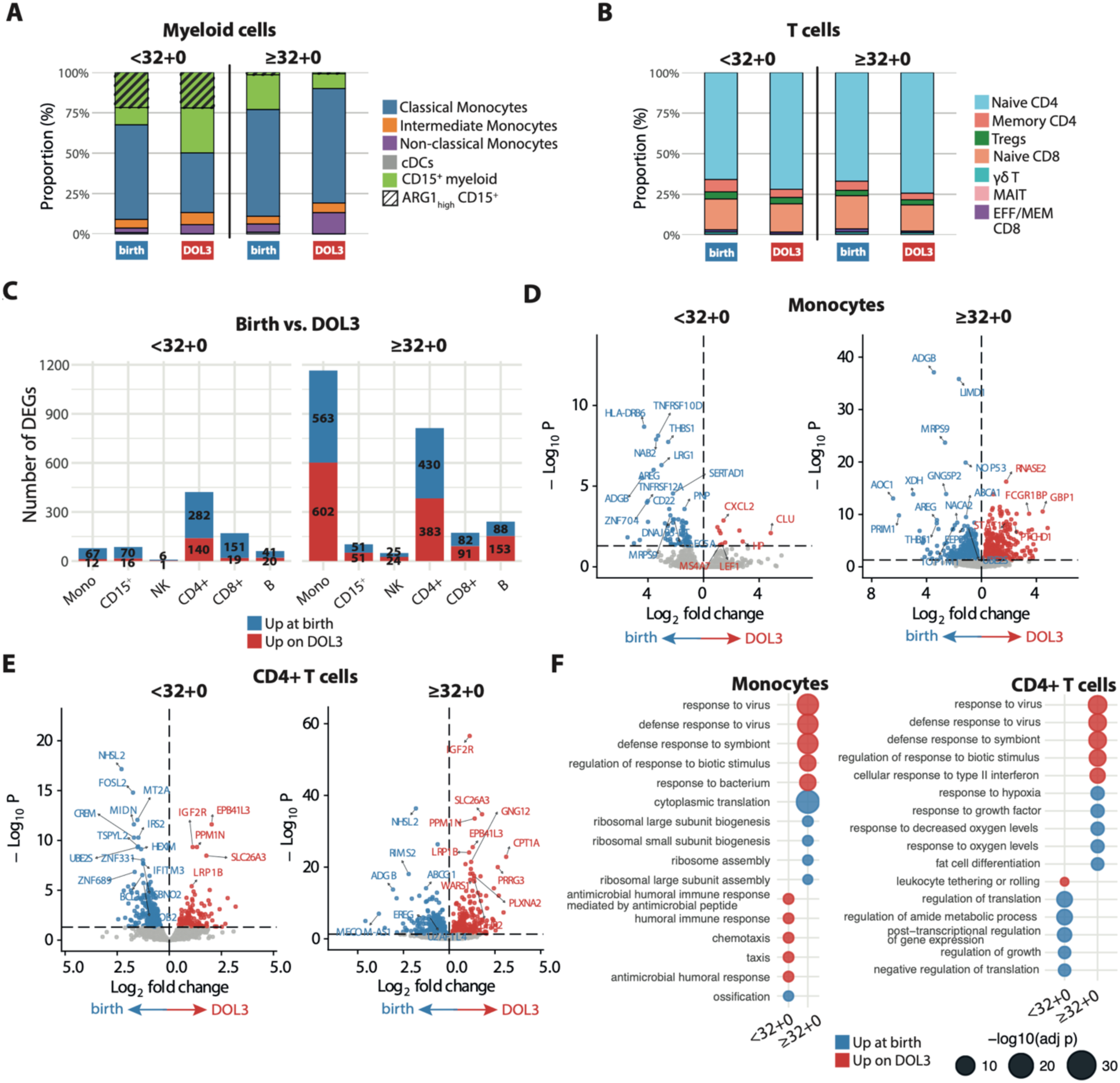
Immune cells from ≥32+0 neonates show enrichment of interferon response pathways on DOL3 at baseline. (A and B) Stacked bar graph representing relative proportion of cells per cell type and group among myeloid cells (A) and T cells (B) at baseline. (C) Stacked bar plot of differentially expressed gene counts by cell type (adj. p < 0.05). (D and E) Volcano plots of baseline gene expression in neonatal monocytes (birth vs. DOL3, D) and CD4+ T cells (birth vs. DOL3, E), stratified by GA-group (adj. p < 0.05). Top DEGs (|log2FC| > 1) were labeled based on ranked adjusted p values. (F) Dot plot of enriched GO Biological Process terms in monocytes (left) and CD4+ T cells (right) for neonates ≥32+0-wk and <32+-wk, based on genes upregulated at birth or on DOL3 (adj. p < 0.05), respectively, with the top 5 terms per condition ranked by adjusted p value (adj. p < 0.05).

To further characterize early postnatal immune changes, we next performed DEG analyses comparing birth and DOL3 samples at baseline. Cells from ≥32-wk neonates exhibited a substantially higher number of upregulated DEGs compared to <32-wk neonates, with monocytes and CD4+ T cells showing the greatest transcriptional changes between birth and DOL3 (Figure 3C). In ≥32-wk monocytes, multiple ISGs, including *GBP1* and *STAT1,* were significantly upregulated on DOL3 relative to birth, and CD4+ T cells also showed induction of ISGs such as *WARS1* (Figures 3D and 3E, Table S17). Monocytes from <32+0-wk exhibited fewer upregulated genes, with key MHC class II genes, *HLA-DRB6* and *HLA-DQA2*, strongly downregulated relative to birth, suggesting reduced antigen-presenting capacity at this time point (Figure 3D, Table S17). Transcriptional upregulation of various ISGs on DOL3 in ≥32-wk neonates was not restricted to monocytes and CD4+ T cells but also seen in CD15⁺ myeloid cells (e.g. *GBP1, IRF1*), CD8+ T cells (e.g. *STAT1, GBP1*), and B cells (e.g. *GBP1, GBP4*). By contrast, cells from <32+0-wk did not show a comparable induction of ISGs across these cell types (Figures S3C, S3E-G, Table S17). GSEA confirmed enrichment of interferon- and antiviral-related programs in ≥32+0-wk, including response to type II interferon and defense response to virus across multiple cell types. In <32+0-wk, no comparable enrichment of antiviral pathways was observed on DOL3, except for monocytes, which instead displayed chemotaxis- and humoral immune response-related enrichment (Figures 3F, S3D, and S3H).

Together, these findings indicate more coordinated interferon-associated transcriptional remodeling by DOL3 in more mature neonates ≥32+0-wk, whereas extremely and very preterm neonates <32+0-wk exhibit larger shifts in cell composition - particularly increased proportions of CD15^+^ myeloid cells, including retained ARG1_high_ CD15^+^ gMDSC-like populations - but comparatively limited transcriptional remodeling.

### GA differentially biases myeloid TLR responses toward interferon-driven vs. KRAS/mTORC1-linked regulatory and metabolic myeloid programs

Building on our observation that GA strongly shapes baseline immune states in neonates, we next asked whether it also affects the neonatal immune cell response to *in vitro* stimulation. To address this, we generated CITE-seq profiles of neonatal immune cells stimulated for two hours with either TLR4 ligand Lipopolysaccharide (LPS) or the TLR7/8 ligand Resiquimod (R848). PCA of the transcriptome data revealed that GA and stimulatory condition, rather than postnatal age were the primary drivers of sample separation (Figure 4A). Consequently, birth and DOL3 samples were jointly analyzed for subsequent analyses to examine GA-dependent responses. Differential expression analysis showed that across cell types, monocytes and CD15^+^ myeloid cells exhibited the strongest transcriptional response following either R848 or LPS stimulation (Figure 4B), in line with their respective TLR expression patterns (*TLR4*, *TLR7*, and *TLR8*; Figure S4A). Interestingly, while baseline transcriptional activity of <32+0-wk samples from extremely and very preterm was modest (Figure 3C), innate stimulation induced differential gene expression that was largely comparable to ≥32+0-wk samples from more mature neonates across all cell types indicating that <32+0-wk immune cells are not dormant but highly reactive upon *in vitro* stimulation. We next focused on myeloid cells to examine their GA-dependent activation in more detail.

**Figure 4.**
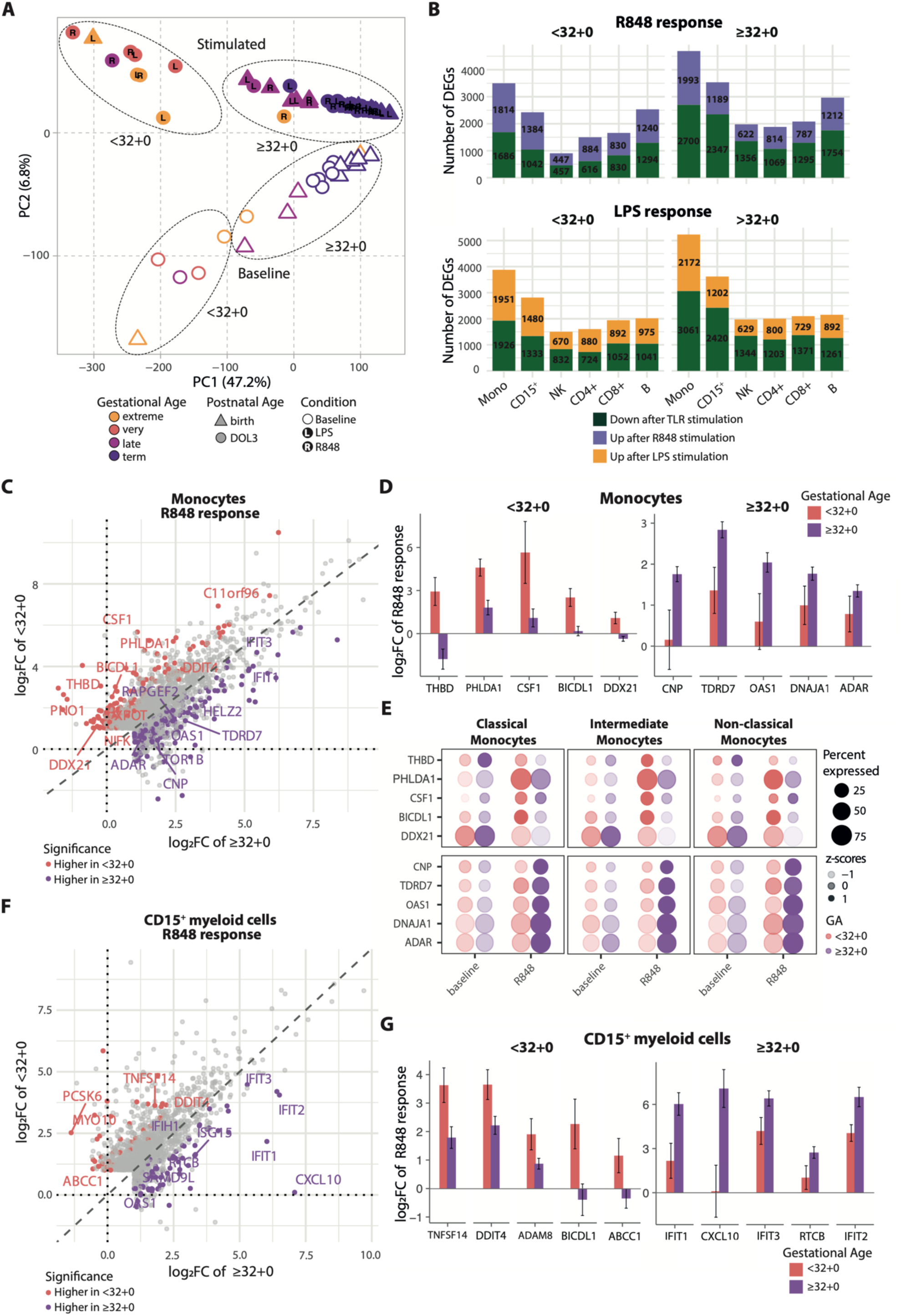
GA differentially biases myeloid TLR responses toward interferon-driven vs. KRAS/mTORC1-linked regulatory and metabolic programs. (A) PCA plot of normalized RNA expression, aggregated by donor and condition. (B) Stacked bar plot of differentially expressed gene counts by cell type (|log₂FC| > 1, adj. p < 0.05, top: R848-stimulated vs. baseline, bottom: LPS-stimulated vs. baseline). (C) Scatter plot of monocyte log₂ fold changes (R848 vs. baseline) for each gene in <32+0-wk (y-axis) and ≥32+0-wk (x-axis) neonates. Genes showing significant interaction effect (adj. p < 0.1) between gestational age and condition are colored. Gene labels indicate the top genes among these interaction-significant genes, ranked by adj. p. (D) Bar plots of log₂ fold changes of the top five genes per age group with significant GA-dependent responses to R848 stimulation in monocytes. Error bars indicate standard errors of the DESeq2 coefficients (lfcSE) for <32+0-wk and ≥32+0-wk neonates following R848 stimulation. (E) Dot plot displaying genes shown in panel D stratified by monocyte subtype. (F) Scatter plot of CD15^+^ myeloid cell log₂ fold changes (R848 vs. baseline), analogous to (C) for monocytes. (G) Bar plots of log₂ fold changes of the top five genes per age group with significant GA-dependent responses to R848 stimulation in CD15^+^ myeloid cells, analogous to (D) for monocytes.

Upon stimulation, <32+0-wk and ≥32+0-wk monocytes and CD15^+^ myeloid cells showed strong induction of multiple inflammatory cytokine and chemokine transcripts, e.g. *IL6, IL1A, CSF3, CCL7,* and *CCL2* (Figures S4B-S4E). Gene set enrichment analyses revealed that both <32+0-wk and ≥32+0-wk activated canonical inflammatory pathways following TLR stimulation, including TNFα, interferon-γ, and IL6/JAK/STAT signaling. Nevertheless, distinct pathways emerged depending on GA. While ≥32+0-wk exhibited stronger enrichment of e.g. interferon-α-driven programs, particularly following LPS stimulation, <32+0-wk showed comparatively enhanced activation of KRAS- and mTORC1-associated signaling pathways (Figures S4F and S4G).

To systematically test for differential responses between the two GA groups, we performed an interaction analysis modeling gene expression as a function of GA group, stimulation, and their interaction (GA group × stimulation). Significance of the interaction term was used to identify genes with GA group-specific stimulation effects. In monocytes, this analysis revealed 338 and 264 differential response genes upon R848 and LPS stimulation, respectively (Figures 4C and S4H, Table S18). Among the genes showing the strongest differential response was thrombomodulin (*THBD*) showing TLR-induced preferential upregulation in <32+0-wk (Figures 4D and S4H). Given the established role of thrombomodulin in modulating inflammatory signaling, promoting monocyte-to-macrophage differentiation, and dampening pro-inflammatory responses, this pattern suggests a bias toward anti-inflammatory and differentiation-associated programs in <32+0-wk monocytes.^30,31^ Notably, these effects were subset-specific: upon R848 stimulation, *THBD* was markedly downregulated in ≥32+0-wk classical monocytes, whereas it was preferentially upregulated in <32+0-wk intermediate monocytes. *CSF1* was likewise strongly induced in the <32+0-wk group. This induction was most pronounced in intermediate monocytes from <32+0-wk preterm infants (Figure 4E). Together, this may point to a potentially self-reinforcing tolerogenic differentiation program within <32+0-wk monocytes. In contrast, ISGs such as *OAS1, IFIT1*, and *IFIT3* were more strongly induced in ≥32+0-wk neonates, particularly within classical and non-classical subsets, consistent with a more antiviral-like response profile (Figures 4C and 4D).

In CD15^+^ myeloid cells, *TNFSF14* (LIGHT) and *DDIT4* emerged as the most strongly GA-dependent genes, preferentially upregulated upon stimulation in <32+0-wk. Whereas *TNFSF14* has been implicated in restraining neutrophil metabolism via LTβR signaling,^32^ *DDIT4* is an mTOR inhibitor induced downstream of IL-10 signaling and has been shown to promote metabolic reprogramming toward oxidative phosphorylation while restraining glycolysis, a shift associated with reduced pro-inflammatory myeloid activation states.^33,34^ Conversely, in ≥32+0-wk CD15^+^ myeloid cells, several ISGs including *IFIT1*, *CXCL10, IFIT3,* and *IFIT2* were among the most strongly differentially regulated genes following R848 stimulation (Figures 4F and 4G). As *CXCL10* is a key chemokine involved in the recruitment of activated T cells and other CXCR3-expressing lymphocytes, its pronounced induction suggests an increased capacity for coordinating downstream adaptive and innate immune responses in ≥32+0-wk neonates.^35,36^ Notably, similar GA-dependent patterns were observed following LPS stimulation, with <32+0-wk samples preferentially upregulating regulatory and metabolic genes such as *THBD* and *TNFSF14*, whereas ≥32+0-wk samples showed stronger induction of canonical ISGs, including *IFIT1, IFIT3, OAS1,* and *MX1* (Figures S4H-S4L).

Collectively, these findings indicate that while both GA groups mount robust transcriptional responses to TLR stimulation, the qualitative nature of these responses differs substantially. Whereas more mature neonates ≥32+0-wk preferentially engage interferon-driven programs, extremely and very preterm neonates <32+0-wk display a shift toward KRAS/mTORC1-linked regulatory and metabolic myeloid programs, suggesting that GA may shape not only cell-intrinsic activation but also downstream immune communication.

### GA shapes predicted T- and NK-cell interactions via myeloid innate cytokine and chemokine signaling

Neonatal T and NK cells showed strong activity after bulk TLR stimulation of CBMC/PBMC samples despite lacking *TLR4*, *TLR7*, and *TLR8* expression, indicating bystander activation via myeloid-derived signals (Figures 4B and S4A). To investigate whether GA influences this intercellular communication, we inferred cell-cell-interactions (CCIs) using CellPhoneDB^37^ restricting the analysis to genes showing significant GA-dependent responses in <32+0-wk or ≥32+0-wk monocytes and CD15^+^ myeloid cells, respectively following R848 stimulation (Figures 5A and 5B).

**Figure 5.**
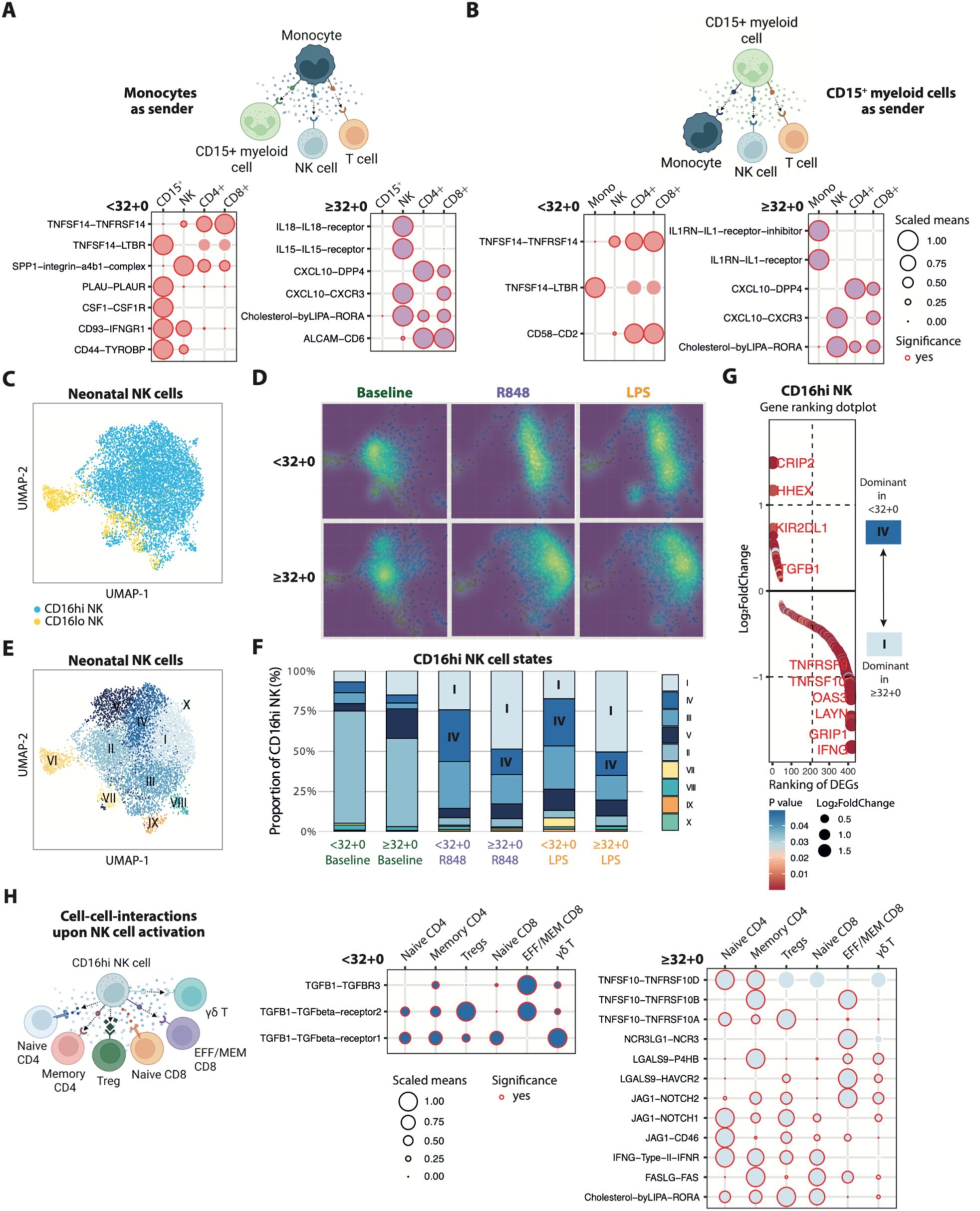
Inferred NK-T cell communication suggests a shift from TGF-β-mediated signaling in <32+0-wk to IFNγ-associated signaling in ≥32+0-wk neonates. (A and B) Top: Schematic of monocyte - CD15^+^ myeloid / NK / T cell - interaction (A). Schematic of CD15^+^ myeloid cell-monocyte / NK / T cell-interaction, created in BioRender (B). Bottom: CellPhoneDB-based dot plots showing ligand-receptor-mediated cell-cell interactions (CCIs) following R848 stimulation, inferred from genes showing significant interaction effects (adj. p < 0.1) between gestational age and condition upregulated in <32+0-wk or ≥32+0-wk neonatal monocytes (A) or CD15^+^ myeloid cells (B). (C) UMAP plot of neonatal NK cells at baseline and after 2 hours stimulation with LPS or R848, derived from neonatal birth and DOL3 samples. (D) Normalized density UMAPs of NK cells split by condition and age group. (E) UMAP of neonatal NK cells at baseline and after 2 hours stimulation with LPS or R848, derived from neonatal birth and DOL3 samples after unsupervised clustering. (F) Stacked bar graph representing relative proportion of cell states among CD16hi NK cells. (G) Gene ranking plot of differentially expressed genes between Cluster IV and Cluster I CD16hi NK cells (adj. p < 0.05), ranked by log₂fold change. (H) Left: Schematic of NK cell - T cell - interaction, created in BioRender. Right: CellPhoneDB-based dot plots showing ligand-receptor-mediated CCIs, inferred from genes enriched in CD16hi NK cell clusters that were preferentially associated with either ≥32+0-wk (cluster I) or <32+0-wk (cluster IV) neonatal samples following TLR stimulation (adj. p < 0.05).

Monocytes exhibited a marked GA-dependent polarization of predicted interaction patterns (Figure 5A). In extremely and very preterm neonates <32+0-wk, monocyte-derived signaling was largely directed toward CD15^+^ myeloid cells. This included PLAU-PLAUR interactions, with PLAUR-expressing neutrophils previously associated with immunosuppressive phenotypes,^38^ as well as TNFSF14-LTBR signaling toward CD15⁺ myeloid cells, consistent with pathways implicated in restraining neutrophil metabolism.^32^ In contrast, monocytes from more mature neonates ≥32+0-wk preferentially engaged lymphoid populations. Predicted interactions were enriched for innate cytokine-mediated signaling, including IL15- and IL18-mediated interactions toward NK cells, alongside CXCL10-CXCR3 signaling targeting NK and CD8+ T cells, indicative of enhanced T- and NK-cell activation.^39,40^

CD15⁺ myeloid cells displayed interaction profiles engaging both myeloid and lymphoid compartments across GA, albeit with distinct emphases (Figure 5B). In <32+0-wk, this included co-stimulatory CD58-CD2 interactions with CD4+ and CD8+ T cells, as well as TNFSF14-mediated signaling toward monocytes. In ≥32+0-wk neonates, CD15⁺ myeloid cells were characterized by anti-inflammatory IL1RN-IL1-receptor signaling toward monocytes and CXCL10-mediated communication with T and NK cells. This included pro-inflammatory CXCL10-CXCR3 interactions with CD8+ T cells and NK cells, together with rather regulatory CXCL10-DPP4 signaling toward CD4+ and CD8+ T cells.^39^

Together, these data indicate that GA shapes predicted immune communication in a cell-type-specific manner. While CD15⁺ myeloid cells interact with both myeloid and lymphoid compartments across gestation, monocytes exhibit a clearer developmental switch around 32 weeks from predominantly myeloid-centered signaling in extremely and very preterm neonates <32+0-wk to lymphoid-directed communication in more mature neonates ≥32+0-wk.

### CD16hi NK cells from ≥32+0-wk neonates exhibit enhanced IFN-γ expression upon R848 stimulation

Predicted CCIs between classical monocytes and NK cells upon stimulation in more mature neonates ≥32+0-wk were enriched for innate cytokine-mediated signaling, including IL-15 and IL-18, both well-known positive regulators of NK cell function.^40,41^ To determine whether NK cell responses also vary with GA, we next focused our analysis on NK cells. Subclustering of neonatal NK cells both at baseline and after bulk TLR stimulation, revealed that CD16hi NK cells diverged by GA under bystander stimulation (Figures 5C and 5D). Within the stimulated CD16hi compartment, NK cluster IV was the dominant bystander state in extremely and very preterm neonates <32+0-wk, whereas NK cluster I was proportionally enriched in more mature neonates ≥32+0-wk (Figures 5E and 5F). DEGs between these clusters were ranked by average log_2_ fold change (Figure 5G). In NK cluster I, leading genes included *IFNG* and *TNFSF10* (encoding TRAIL), whereas *KIR2DL1* (encoding an inhibitory NK cell receptor) ranked among the top three genes in NK cluster IV. Hierarchical clustering of these DEGs further highlighted distinct age-dependent patterns among CD16hi NK cells (Figures S5A and S5B): in ≥32+0-wk, bulk TLR stimulation strongly upregulated ISGs as well as genes associated with NK cell effector function such as *IFNG, GNLY, GZMB, PRF1, FASLG*, and *TNFSF10*. Genes involved in innate cytokine signaling, such as *IL15RA, IL12RB1, IL18R1*, and *RUNX3*^42^ were also upregulated in ≥32-wk CD16hi NK cells. By contrast, in <32+0-wk, several apoptosis-associated genes, the inhibitory receptor *KIR2DL1* and *TGFB1,* which has been shown to suppress NK cell proliferation and activation,^43,44^ were preferentially expressed upon stimulation.

Overall, these results indicate that CD16hi NK cells also exhibit GA-dependent divergence upon bulk TLR stimulation, with CD16hi NK cells from more mature neonates ≥32+0-wk exhibiting enhanced IFN-γ expression upon R848 stimulation.

### Predicted NK-T cell crosstalk shifts from TGF-β-dominated regulation in <32+0-wk to concurrent activation and regulatory signaling via IFN-γ and TRAIL in ≥32+0-wk

NK cells can shape early T cell priming and Th polarization both directly and indirectly through antigen-presenting cells (APCs). In particular, NK cell-derived IFN-γ promotes the polarization of naive CD4+ T cells toward Th1 cells,^45,46^ while TGF-β has been shown to promote the differentiation into Foxp3+ regulatory T cells (Tregs),^46–48^ Building on the observed GA-dependent differences in TLR-induced NK bystander responses, we next examined whether the predicted CCIs between CD16hi NK cells and T cells also vary with GA. To address this, ligand-receptor interactions were inferred using genes enriched in CD16hi NK cell clusters that were proportionally dominant in either ≥32+0-wk (NK cluster I) or <32+0-wk neonatal samples (NK cluster IV).

In extremely and very preterm neonates <32+0-wk, predicted NK-T cell interactions were dominated by TGF-β-mediated signaling. In contrast, NK-T cell crosstalk in more mature neonates ≥32+0-wk was characterized by enriched IFNG-mediated signaling toward multiple T cell subsets, together with TRAIL-and FASLG-mediated interactions involving both pro-apoptotic (e.g. TNFRSF10A, FAS) and anti-apoptotic effects (e.g. TNFRSF10D), depending on the repertoire of receptors expressed by the interacting T cells (Figure 5H).

These CCIs engaged nearly all T cell subsets, suggesting that NK-T cell crosstalk in ≥32+0-wk neonates integrates both activating and regulatory axes, potentially establishing a feedback loop that tempers excessive T cell activation. By contrast, TGF-β-dominated NK-T cell crosstalk in <32+0-wk neonates may reflect a developmentally imposed regulatory state limiting Th1 polarization.

### Gestational age shapes Treg- and Th1-associated transcriptional programs in neonatal CD4+ naïve T cells upon bystander activation

Given the observed GA-dependent differences in NK-T cell crosstalk, we next asked whether these cues translated into divergent transcriptional programs within neonatal T cells upon bystander activation. Clustering of neonatal T cells at both baseline and after TLR stimulation, revealed that particularly CD4+ naïve T cell states diverged by GA during bystander activation (Figures 6A and 6B). Because multiple innate immune cell types contribute to the cytokine milieu during bystander stimulation, we next used cytokine activity inference^49^ to identify innate-derived cytokines most strongly associated with, and potentially modulating, GA-dependent T cell responses. Cytokine activity inference showed that naïve CD4+ T cells from more mature neonates ≥32+0-wk preferentially upregulated e.g., IFN-γ-, IL-12-, and IL-15-associated signatures upon both R848 and LPS stimulation, consistent with early Th1 polarization (Figures 6C, S6A, and S6B).^50,51^ In contrast, naïve CD4+ T cells from extremely and very preterm neonates <32+0-wk exhibited e.g. enriched IL-10 signaling following R848 stimulation and enriched IL-1 receptor antagonist-associated signatures under both stimulatory conditions, suggesting a more regulatory immune milieu,^52,53^ with IL-16-associated signaling also enriched in this group. In addition, Tregs from ≥32+0-wk neonates showed enrichment of GITRL-mediated signaling upon R848 stimulation, consistent with attenuated Treg suppressive activity and enhanced immune activation.^54,55^

**Figure 6.**
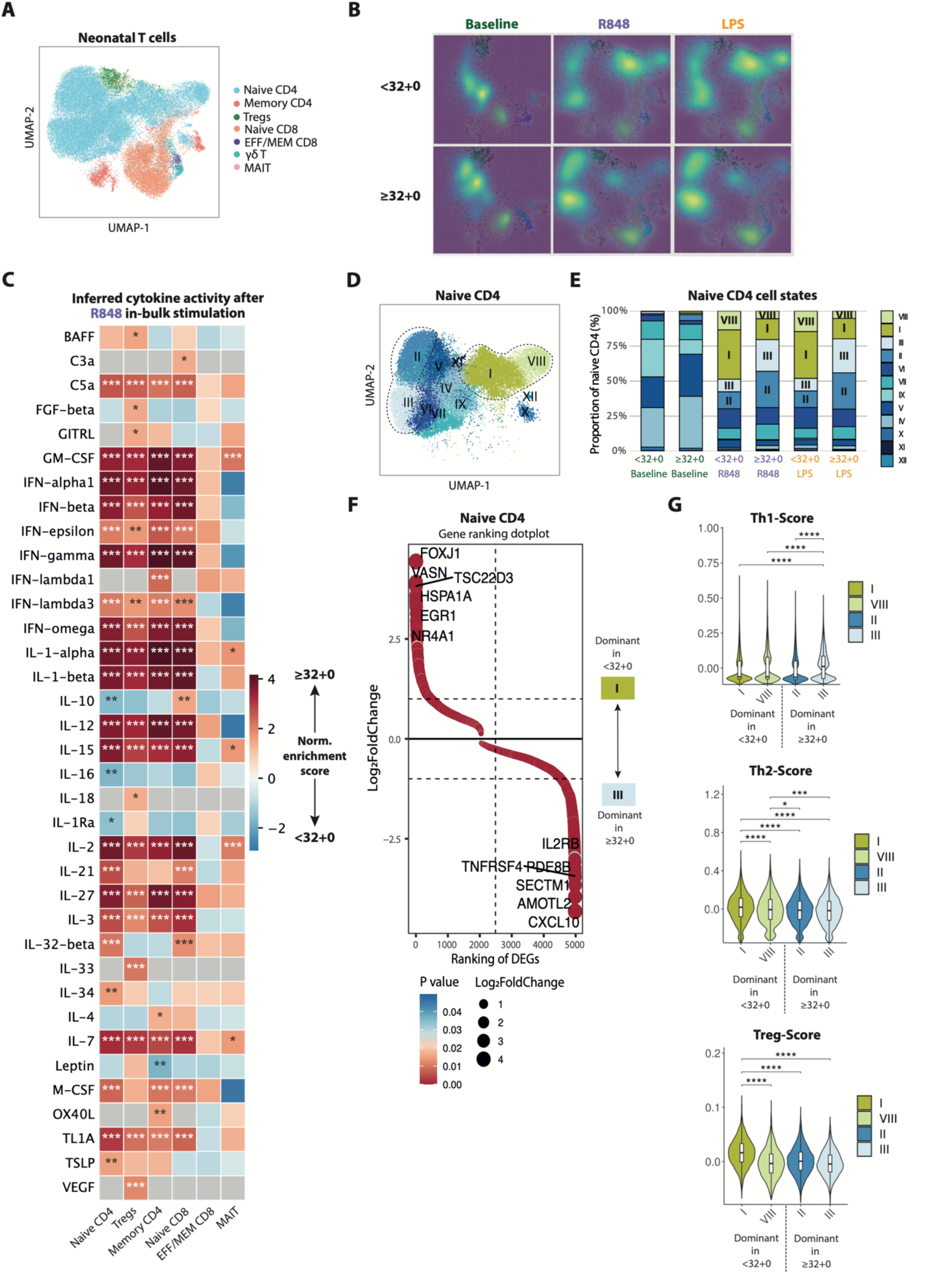
Naïve CD4+ T cells from <32-wk neonates preferentially engage regulatory programs, whereas ≥32-wk neonates display enhanced Th1-associated activation upon bystander activation. (A) UMAP plot of neonatal T cells at baseline and after 2 hours stimulation with LPS or R848, derived from neonatal birth and DOL3 samples. (B) Normalized density UMAPs of T cells split by condition and age group. (C) Heatmap showing results from huCIRA-based gene set enrichment analysis of cytokine-associated gene signatures in T cell subsets comparing ≥32+0 and <32+0 neonatal samples after R848 stimulation.^49^ Cytokines are shown if significant enrichment was observed in at least one T cell subset. Red: enrichment in ≥32+0-wk, blue: enrichment in <32+0-wk samples. Asterisks denote statistically significant enrichment. (D) UMAP of neonatal naive CD4+ T cells at baseline and after 2 hours stimulation with LPS or R848, derived from neonatal birth and DOL3 samples after unsupervised clustering. (E) Stacked bar graph representing relative proportion of cell states among naïve CD4+ T cells. (F) Gene ranking plot of differentially expressed genes between Cluster I and Cluster III naïve CD4 T cells (adj. p < 0.05), ranked by log₂ fold change. (G) Violin plots show gene module scores for Th1-, Th2-, and Treg-associated gene sets across naïve CD4 T cell states (calculated at the single-cell level; pairwise Wilcoxon rank-sum tests; adj p: * < 0.05, ** < 0.01, *** < 0.001, **** < 0.0001). Effect sizes were estimated using Cliff’s delta and are reported alongside adjusted p values in Table S12.

Focusing specifically on naïve CD4+ T cell heterogeneity, subclustering revealed that CD4 clusters I and VIII predominated in <32+0-wk neonates, whereas CD4 clusters II and III were enriched in ≥32+0-wk neonates (Figures 6D and 6E). DGE analysis between the dominant clusters (I vs. III and VIII vs. III) further highlighted distinct transcriptional programs (Figures 6F and S6C).

CD4 cluster I, dominant in <32+0-wk neonates, was characterized by elevated expression of *FOXJ1, VASN*, *EGR1*, and *NR4A1. FOXJ1* and *NR4A1* are known negative regulators of T cell activation and effector programming, while *EGR1* preferentially supports Th2-associated transcriptional responses, and *VASN* has been linked to suppression of T cell activation via AKT-dependent signaling.^56–59^

CD4 cluster VIII, also proportionally enriched in <32+0-wk neonates, expressed higher levels of *SNAI1, TIGIT, FOXP3,* and *TXNIP*, consistent with regulatory, suppressive, or contraction-associated T cell states.^60–63^ In contrast, CD4 cluster III, enriched in ≥32+0-wk neonates, showed increased expression of *CXCL10*, *AMOTL2, TNFRSF4* (OX40)*, PDE8B, IL2RB*, and *BHLHE40* reflecting enhanced interferon responsiveness, migratory capacity, co-stimulatory signaling, and pro-inflammatory effector programming ^64–69^. Consistent with these transcriptional differences, gene module scoring based on published Th1-, Th2-and Treg-associated signatures^9,70^ revealed that <32+0-wk-dominant CD4 clusters exhibited higher Treg-associated and modest Th2-associated scores, whereas ≥32+0-wk CD4 cluster III showed increased Th1-associated signatures (Figure 6G). Notably, this tolerance-associated bias in <32+0-wk neonates was supported across independent analytical layers, including cytokine activity inference (IL-10 and IL-1RA-associated signaling), cluster-defining gene expression, and Treg module scoring.

Together, these findings indicate that GA critically shapes the balance between regulatory and effector-associated transcriptional programs of neonatal CD4+ T cells upon bystander activation. While more mature neonates ≥32+0-wk exhibited a Th1-biased transcriptional response aligned with the IFN-γ-dominated NK-T cell crosstalk, extremely and very preterm neonates <32+0-wk displayed enrichment of Treg-associated programs consistent with the broader regulatory immune milieu characterized by CD15^+^ myeloid cells exhibiting gMDSC-like features and the enhanced TGF-β-mediated NK-T cell interactions.

## DISCUSSION

In this study, we present, to our knowledge, the first comprehensive neonatal immune CITE-seq atlas capturing the first days of life across a wide range of gestational ages (GA), integrating both baseline immune states and responses to TLR stimulation. By combining multimodal profiling with functional perturbation, we identify a definable developmental transition around ∼32 weeks of gestation. This threshold separates two distinct immunobiological states: a ‘CD15^+^ myeloid gMDSC-dominated’ phenotype in extremely and very preterm infants and an ‘interferon-primed’ phenotype in late preterm and term infants, resulting in divergent myeloid-to-lymphocyte signaling and qualitatively distinct NK- and T-cell bystander responses upon innate stimulation. Together, these findings support a model of layered immune ontogeny, in which developmental programs - most notably the establishment of tonic interferon activity and the contraction of suppressive myeloid populations - may act as gating events in immune maturation.^12,15,71^ Our data suggest that failure to initiate these programs may fundamentally alter how the neonatal immune system perceives and responds to early postnatal inflammatory cues, thereby contributing to the heightened immune vulnerability observed in extremely and very preterm infants.

At baseline, we observed that the neonatal immune system compared to the adult is not quiescent but exists in a gestational age-dependent fine-tuned state characterized by the coexistence of regulatory features and broad signs of basal preactivation. In neonates born at or after 32 weeks, this state was defined by a rapid upregulation of interferon response signatures by the third day of life. This aligns with previous multi-omic analyses of healthy term newborns, which described a robust developmental trajectory dominated by interferon signaling during the first week of life.^15^ Furthermore, this upregulation likely represents the successful initiation of an ISG-high baseline, a constitutive antiviral state recently described in healthy infants that may poise the immune system for rapid pathogen defense.^16,72^ Crucially, our data showed that extremely and very preterm infants born before 32 weeks do not initiate this program immediately after birth. While the precise ontological drivers of this transition in humans remain to be fully elucidated, our findings parallel recent mechanistic findings from murine models. Li et al. have demonstrated that a sterile, skin-derived ‘interferon spike’ occurs specifically during late gestation, acting as a developmental pulse that forces hematopoietic stem (HSCs) and progenitor cells (HPCs) to transition from fetal to adult transcriptional states in mice.^73^ Our findings suggest that extremely and very preterm human infants may be delivered before this critical switch occurs. Consequently, they might enter the extrauterine environment in a relative ‘interferon void’, missing the physiological cue required to mature their immune trajectory on time. While recent longitudinal studies indicate that extremely premature infants eventually develop high interferon signatures by two months of life,^9^ our data imply that the absence of this program in the immediate postnatal period may leave these infants in a vulnerable, distinct state driven by compositional rather than transcriptional adaptation.

The immune landscape of extremely and very preterm neonates <32+0-wk was dominated by cell subsets with potentially suppressive or regulatory properties, particularly CD15^+^ myeloid cells with a gMDSC-like (granulocytic myeloid-derived suppressor cell-like) phenotype. The exact frequency and functionality of gMDSCs in preterm infants remains controversial. While some studies report reduced frequencies of gMDSCs with diminished suppressive function in preterm or very low birth weight infants compared to term controls,^13,74^ others describe increased gMDSC proportions in cord blood independent of gestational age, persisting in the peripheral blood of preterm infants throughout the neonatal period.^75^ These discrepancies likely reflect technical and conceptual challenges in defining this heterogenous population, particularly when relying on unimodal approaches such as flow cytometry or transcriptomics alone. A key strength of our multimodal CITE-seq approach lies in the simultaneous integration of surface protein phenotypes (e.g., CD15, CD16 surface expression) with intracellular transcriptional programs (e.g., *ARG1* expression) at the single-cell resolution. This integrated analysis enabled the identification of ARG1_high_ CD15^+^ gMDSC-like myeloid cells that were preferentially enriched in <32+0-wk infants. Notably, this enrichment occurred in parallel with reduced interferon-associated transcriptional signatures, suggesting a potential mechanistic link. Type I interferon signaling has been described as a critical negative regulator of MDSC differentiation.^76^ Specifically, the absence of interferon signaling has been shown to release the constraint on the PI3K-Akt-mTOR pathway, a key metabolic driver of *ARG1* expression and suppressive function.^77^ Our data, which reveal a concurrence of low interferon signatures, proportional enrichment of ARG1_high_ CD15^+^ gMDSC-like myeloid cells, and enriched mTORC1 signaling in the myeloid compartment of <32+0-wk infants, support this model. Furthermore, the expansion of ARG1_high_ CD15^+^ myeloid cells may reflect a dysregulated attempt at emergency myelopoiesis. Recent evidence suggests that fetal hematopoietic progenitors are restricted from engaging in emergency myelopoiesis by maternal IL-10 to prevent immunopathology *in utero.*^78^ Preterm birth may abruptly release this maternal restraint in a host with a progenitor pool that is developmentally biased toward regulatory myeloid differentiation,^79^ potentially triggering a dysregulated form of myelopoiesis that preferentially expands ARG1_high_ gMDSC-like myeloid cells.

Upon *in vitro* TLR stimulation, myeloid cells from both GA groups mounted robust responses, yet the qualitative nature of these responses diverged. While cells from more mature infants ≥32+0 preferentially activated interferon-driven antiviral programs, those from <32+0-wk infants exhibited responses enriched for KRAS- and mTORC1-associated signaling and regulatory pathways. This divergence further extended to downstream lymphocyte bystander activation. Despite minimal expression of *TLR4*, *TLR7*, and *TLR8* in neonatal T and NK cells, both cell types exhibited strong activation, confirming recent evidence that neonatal lymphocytes are highly sensitive to TCR-independent bystander stimulation.^21^ However, the outcome of this activation was highly GA-dependent. CD16hi NK cells from ≥32+0-wk neonates displayed enhanced IFN-γ expression and predicted NK-T cell crosstalk involved both activating and regulatory cues, potentially preventing excessive activation. In contrast, inferred NK-T cell communication in <32+0-wk neonates was dominated by TGF-β-mediated regulatory signaling. This regulatory bias extended to the adaptive compartment, where naïve CD4+ T cells preferentially adopted Treg-associated transcriptional programs in the <32+0-wk group, consistent with the fetal predisposition toward tolerance described in layered immune ontogeny,^80^ whereas ≥32+0-wk T cells shifted toward Th1-effector-associated states upon activation. Collectively, these findings indicate that the distinct immune state of extremely and very preterm neonates <32+0-wk is not a result of functional immaturity, but may rather reflect the retainment of a fetal developmental layer optimized for *in utero* tolerance.^3,80^ While this program serves to prevent immunopathology during development, its persistence ex utero allows extremely and very preterm infants to respond to stimulation while remaining developmentally constrained in mounting a canonical effector response.

Overall, our findings identify ∼32 weeks of gestation as a key developmental transition in neonatal immune maturation that coincides with a broader clinically relevant phase, during which outcomes of prematurity markedly improve. This period also aligns with maturation of other organ systems, including the establishment of sufficient pulmonary surfactant production, a process also influenced by inflammatory signaling pathways, suggesting coordinated immune-tissue development. The emergence of tonic interferon signaling and the reduction of suppressive myeloid programs suggest a coordinated transition toward an immune state prepared for the challenges of postnatal transition and extrauterine life. Failure to initiate this transition may leave extremely and very preterm infants in a fetal-programmed immune state that favors regulation over effector responses, potentially predisposing to dysregulated immune adaptation and later inflammatory injury. While these findings raise the possibility that aspects of immune maturation could be therapeutically supported, they also highlight important constraints: excessive interferon activity during development may carry risks, and suppressive myeloid cells were reported to serve protective roles in early life.^13^ Any attempt to modulate these pathways will therefore require careful consideration of developmental timing, tissue readiness, and unintended consequences in preterm infants.

### Limitations of the study

Several limitations should be considered when interpreting these findings. First, our sample size (82 samples from 25 neonates and 10 adults) reflects the inherent technical and ethical challenges of studying human neonatal immunity and may limit statistical power for detecting more subtle effects. In addition, while our data support the presence of an ARG1_high_ CD15^+^ gMDSC-like dominated phenotype in <32+0-wk infants, functional validation is needed to confirm its suppressive capacity and developmental trajectory in early life. *In vitro* TLR stimulation is a simplified model and does not capture the full complexity of neonatal inflammatory exposures *in vivo*. However, it enables direct comparison of intrinsic immune response programs across gestational ages under controlled conditions. Cell-cell interaction analyses are based on computational inference and therefore remain predictive, but they are well suited to identify dominant signaling axes in systems that are difficult to manipulate experimentally. Our analysis is limited to the immediate postnatal period and does not address the long-term stability of the observed immune states; nevertheless, this early window represents a critical phase of immune adaptation preceding many complications of prematurity. Finally, we cannot distinguish to what extend the observed effects are driven by intrinsic programs versus external cues such as antenatal corticosteroid exposure or antibiotic treatment. Because these exposures are tightly linked to the clinical management of extreme prematurity, disentangling their specific contributions is inherently challenging in studying human neonatal immunity, where limited blood volumes constrain extensive functional analyses. Nevertheless, our findings reflect the ‘real-life’ immune state of preterm infants and therefore remain highly relevant for understanding early immune vulnerability.

## RESOURCE AVAILABILITY

### Lead contact

Requests for further information and resources should be directed to and will be fulfilled by the lead contact, Prof. Dr. Sarah Kim-Hellmuth.

### Materials availability

This study did not generate new unique reagents.

### Data and code availability

All data associated with this study are present in the paper, the Supplementary Materials, or as indicated as follows: The CITE-seq dataset from this study will be available for interactive exploration and download through the CELLxGENE web portal. The raw CITE-seq and processed data matrices will be available at Gene Expression Omnibus. Custom scripts used for data analysis and visualization are available at https://github.com/kimhellmuthlab/neociteseq_atlas. All other analyses were performed using publicly available software packages as described in the Methods.

## ACKNOWLEDGEMENTS

We would like to thank all study participants and their families for taking part in this study. We are greatful to the Division of Neonatology and the Clinic for Obstetrics & Gynecology at LMU University Hospital, specifically the clinical study team, all midwifes, and clinicians, for contributions to patient enrollment, sample collection, and support of this study. We would like to thank the Core Facility Genomics at Helmholtz Munich, specifically Sudharsan Padmarasu and Inti De La Rosa Velazquez for library prep and sequencing. We are grateful to the Flow Cytometry Core Facility at the Comprehensive Childhood Research Center of Dr. von Hauner Children’s Hospital, particularly Raffaele Conca and Susanne Wullinger for assistance with flow cytometry analysis and panel design. We would like to thank Juliane Klein for her technical assistance. We are grateful to Eleftheria Zeggini, Sagar, Dorothee Viemann, Jessica Jin, Marie Bourdon, and Florens Lohrmann for valuable discussions and helpful suggestions that improved this study.

P.R., A.M., M.I.C., P.M.-K., M.S., P.H., C.N., and S.K.-H were funded by the German Research Foundation (DFG) within TRR359 (Project ID 491676693). C.N. is further supported by the Medical & Clinician Scientist Program (MCSP) of the medical faculty of the Ludwig-Maximilians-University. S.K.-H. is further supported by the DFG Emmy Noether Programme KI 2091/2-1 (459153572), SFB/TRR237-B29 (369799452), BMBF DZKJ (01GL2406A), Cluster for Nucleic Acid Sciences and Technologies - NUCLEATE (533767322 - EXC 3113/1), and an ERC Starting Grant (101076303).

## AUTHOR CONTRIBUTIONS

Conceptualization, S.K.-H and C.N.; methodology, P.R.; software, B.P. and P.R.; formal analysis, P.R.; investigation, P.R.; validation, P.R., A.M., M.I.C., P.M.K. and J.W.; visualization, P.R., A.M. and M.I.C.; funding acquisition, S.K.-H and C.N.; project administration, S.K.-H and C.N.; supervision, S.K.-H and C.N.; writing – original draft, P.R., S.K.-H and C.N.; writing – review & editing, M.S., P.H. and C.K.

## SUPPLEMENTAL INFORMATION

Document S1. Supplementary Material and Methods, Figures S1 – S6, Tables S1 – S15, and legends for Tables S16 – S18

Table S16. Differentially expressed genes between neonatal and adult immune cells

Table S17. Differentially expressed genes between birth and DOL3

Table S18. Differentially expressed genes between LPS/R848 and baseline

## METHOD DETAILS

### Study design and donor characteristics

Donor characteristics are presented in Tables S1 - S7 and Tables S10 - S11. Patients were enrolled abiding by the Declaration of Helsinki and with LMU University’s institutional review board (IRB) approval (IRB no. 15-0735_1 and 22-0534). All neonates born at the perinatal centers of the LMU Klinikum within predefined gestational age (GA) categories were eligible for inclusion: extremely preterm (<28+0 weeks of gestation), very preterm (28+0 to 31+6 weeks), late preterm (32+0 to 36+6 weeks), and full-term infants (≥37+0 weeks). Exclusion criteria included major congenital anomalies, syndromic disease, and known hematopoietic disorders. Umbilical cord blood samples from late preterm and term infants were collected immediately after delivery as anonymized residual material that would otherwise have been discarded. Therefore, clinical metadata for these donors are not available. For the term infants, cord blood was obtained exclusively from uneventful, planned primary cesarean sections. Peripheral blood samples were obtained between 36 and 72 hours of life during routine newborn screening or at other routine blood draws following written informed consent from the parents or legal guardians. A minimum of 150μl of blood was collected into EDTA-coated Microvette tubes (Sarstedt) and processed within 2 hours of collection. Peripheral blood mononuclear cells (PBMCs) and cord blood mononuclear cells (CBMCs) were isolated by density gradient centrifugation and cryopreserved using an optimized protocol for very low cell counts, as detailed in the Supplementary Materials and Methods.

### In vitro stimulation and AbSeq labeling

After thawing, samples were washed once with pre-warmed complete medium (RPMI supplemented with 10% fetal calf serum and 1% penicillin/streptomycin) to remove cryopreservation media. Cells were resuspended at 1 x 10^6^ cells/mL, pooled by gestational age category and subsequently split into unstimulated and stimulated conditions. *In vitro* stimulation was performed for 2h at 37 °C with lipopolysaccharide (LPS; 200 ng/mL; from *Escherichia coli*, Sigma-Aldrich, L2654; reconstituted in sterile phosphate-buffered saline [PBS]) or the TLR7/8 agonist Resiquimod (R848; 500 ng/mL; InvivoGen; reconstituted in dimethyl sulfoxide [DMSO]). Following stimulation, cells were washed with PBS, centrifuged (400 x g, 10 min), and resuspended at 1 x 10^6^ cells/mL. CITE-seq was then performed according to the manufacturer’s protocol using the BD Rhapsody^TM^ Single-Cell Analysis System. Samples were labeled with SampleTags (BD® Single-Cell Multiplexing Kit) and subsequently labeled with a customized panel of 36 oligonucleotide-conjugated antibodies, consisting of the BD® AbSeq Immune Discovery Panel (30 markers) and six additional antibodies targeting neonatal-enriched immune cell populations (Tables S8 and S9). In total, 400,000 labeled cells were processed for single-cell capture according to the manufacturer’s instructions.

### CITE-seq

Single-cell cDNA with matched Sample Tag and AbSeq libraries were prepared using the BD Rhapsody^TM^ cDNA Kit and the BD Rhapsody^TM^ Whole Transcriptome Analysis Amplification Kit according to the manufacturer’s instructions and sequenced on an Illumina NovaSeq 6000 or NovaSeq X plus platform to obtain paired-end reads.

### CITE-seq data analysis

Raw sequencing data were processed using the BD Rhapsody^TM^ Sequence Analysis Pipeline according to the manufacturer’s instructions to generate gene expression and AbSeq count matrices as well as assigned Sample Tag identities. After this preprocessing pipeline 195,000 cells were identified for further analysis. Donor identities were resolved by genotype-based *in silico* demultiplexing using vireo.^81^ Computational doublet detection was performed using scDblFinder.^82^ 61,260 cells classified as doublets or unassigned by genotype-based demultiplexing, scDblFinder, or Sample Tag assignment were excluded from downstream analyses. To filter out low-quality cells, cells with fewer than 500 detected genes, unusually high gene or UMI counts, or more than 25% mitochondrial reads were excluded from the analysis. After strict quality control filtering 115,037 cells were used for downstream analysis. Multimodal data integration was performed in Python using totalVI.^83^ Leiden clustering was performed using Scanpy.^84^ Principal component analysis (PCA) was performed on donor- and condition-averaged, normalized gene expression profiles using the prcomp function in R. Differences in global gene expression profiles were assessed using permutational multivariate analysis of variance (PERMANOVA; adonis2 function, vegan package) on Euclidean distances of log-normalized data, with 999 permutations. For within-donor comparisons, permutations were stratified by donor identity. Homogeneity of dispersion was assessed using betadisper. Cluster annotation was performed based on totalVI-derived Leiden clusters. Marker genes for each cluster were identified using the Seurat FindAllMarkers workflow,^85^ and clusters were manually annotated on the basis of differentially expressed genes (DEGs) in each cluster and established canonical marker genes. Cell type assignments were further validated using AbSeq-derived surface protein expression profiles. ADT feature plots show totalVI-denoised protein expression. Normalized density UMAP plots were generated by calculating two-dimensional kernel density estimates on UMAP embeddings for defined sample groups and visualized as normalized density overlays. Cell type compositional differences were assessed using the Bayesian compositional analysis framework scCODA.^23^ As no prior knowledge about an invariant cell type was available, the reference cell type was selected automatically by scCODA.

Pseudobulk differential gene expression analysis was performed using DESeq2 on donor- and sample-aggregated cell counts. Interaction models were used exclusively for GA group-specific stimulation effects, whereas stimulated vs. baseline comparisons included batch as a covariate, and adult vs. neonatal and DOL3 vs. birth comparisons were performed using condition-only models. Error bars indicate 95% confidence intervals derived from the DESeq2 standard errors of log_2_ fold change estimates. Following differential gene expression analysis, results were filtered to exclude ENSG-only and LINC-annotated genes, thereby restricting the results to annotated, predominantly protein-coding genes. Heatmaps of scaled average gene expression were generated using the pheatmap package with hierarchical clustering applied to rows. Dot plots of DEGs were generated using the DotPlot2 function from the SeuratExtend package in R.^86^ Furthermore, DEGs were visualized by volcano plots using the R package EnhancedVolcano. Gene ranking plots were generated by ranking DEGs by log_2_ fold change and visualized using the gene_rank_plot function from the TOmicsVis package in R.^87^ UpSet plots were generated using the ComplexUpset package in R to visualize intersections of gene sets across immune cell types. Gene set enrichment analysis of single-cell-derived DEG lists was performed using Enrichr with the MSigDB Hallmark 2020 gene set collection.^88^ For a selected analysis, Gene Ontology Biological Process enrichment was performed using clusterProfiler with the DESeq2-tested genes used as background.^89^

Cell-cell interactions (CCIs) were inferred using CellPhoneDB^37^ employing the DEG-based analysis mode. For monocytes and CD15^+^ myeloid cells, ligand-receptor interactions were analyzed using genes identified from the GA group-specific stimulation interaction analysis (GA group × stimulation, adj. p < 0.1) following R848 stimulation, analyzed separately for <32+0-wk and ≥32+0-wk samples. In a complementary analysis for NK cells, CCIs were inferred using genes enriched in CD16hi NK cell clusters dominant in either <32+0-wk or ≥32+0-wk samples (adj. p < 0.05). CCI results were visualized using dot plots generated with the R package ktplots.^37^

Cytokine-associated enrichment scores were computed using the huCIRA framework, which applies a preranked GSEA based on differential gene expression between conditions.^49^

Single-cell module scores for Th1, Th2, and Treg gene sets were computed using Seurat’s AddModuleScore on normalized data. Gene sets were derived from published T helper signatures^9^ and the MSigDB C7 immunologic signature collection.^70^ Module score distributions were visualized using enhanced violin plots generated with the Vlnplot2 function from the R package SeuratExtend.^86^

### Multicolor flow phenotyping

Multicolor flow cytometry was performed on a BD LSRFortessa^TM^. Fc receptor blocking was performed prior to antibody staining (BD Pharmingen^TM^ Human BD Fc Block^TM^). Fluorochrome-conjugated antibodies used are listed in Tables S13 - S15. Data were analyzed using FlowJo^TM^ v10.

### Statistical analysis

Statistical analyses for multicolor flow cytometry data were performed using R. Data distributions were assessed for normality using the Shapiro-Wilk Test prior to statistical testing. For non-normally distributed data, pairwise comparisons were performed using Wilcoxon rank-sum tests with Bonferroni correction for multiple testing (pBonf). For normally distributed data, homogeneity of variances was assessed using Levene’s test, followed by one-way analysis of variance (ANOVA) where appropriate. Statistical significance levels are indicated as follows: adj. p: * < 0.05, ** < 0.01, *** < 0.001, **** < 0.0001.

## REFERENCES

1. Kollmann, T.R., Kampmann, B., Mazmanian, S.K., Marchant, A., and Levy, O. (2017). Protecting the Newborn and Young Infant from Infectious Diseases: Lessons from Immune Ontogeny. Immunity 46, 350–363. 10.1016/j.immuni.2017.03.009.

2. Henneke, P., Kierdorf, K., Hall, L.J., Sperandio, M., and Hornef, M. (2021). Perinatal development of innate immune topology. eLife 10, e67793. 10.7554/eLife.67793.

3. Rudd, B.D. (2020). Neonatal T Cells: A Reinterpretation. Annu. Rev. Immunol. 38, 229–247. 10.1146/annurev-immunol-091319-083608.

4. Bennett, N.J., Tabarani, C.M., Bartholoma, N.M., Wang, D., Huang, D., Riddell, S.W., Kiska, D.L., Hingre, R., Rosenberg, H.F., and Domachowske, J.B. (2012). Unrecognized Viral Respiratory Tract Infections in Premature Infants during their Birth Hospitalization: A Prospective Surveillance Study in Two Neonatal Intensive Care Units. J. Pediatr. 161, 814–818.e3. 10.1016/j.jpeds.2012.05.001.

5. Das, A., Ariyakumar, G., Gupta, N., Kamdar, S., Barugahare, A., Deveson-Lucas, D., Gee, S., Costeloe, K., Davey, M.S., Fleming, P., et al. (2024). Identifying immune signatures of sepsis to increase diagnostic accuracy in very preterm babies. Nat. Commun. 15, 388. 10.1038/s41467-023-44387-5.

6. Vatanen, T., Kostic, A.D., d’Hennezel, E., Siljander, H., Franzosa, E.A., Yassour, M., Kolde, R., Vlamakis, H., Arthur, T.D., Hämäläinen, A.-M., et al. (2016). Variation in Microbiome LPS Immunogenicity Contributes to Autoimmunity in Humans. Cell 165, 842–853. 10.1016/j.cell.2016.04.007.

7. Laforest-Lapointe, I., and Arrieta, M.-C. (2017). Patterns of Early-Life Gut Microbial Colonization during Human Immune Development: An Ecological Perspective. Front. Immunol. 8, 788. 10.3389/fimmu.2017.00788.

8. Olin, A., Henckel, E., Chen, Y., Lakshmikanth, T., Pou, C., Mikes, J., Gustafsson, A., Bernhardsson, A.K., Zhang, C., Bohlin, K., et al. (2018). Stereotypic Immune System Development in Newborn Children. Cell 174, 1277–1292.e14. 10.1016/j.cell.2018.06.045.

9. Olaloye, O., Gu, W., Gehlhaar, A., Sabuwala, B., Eke, C.K., Li, Y., Kehoe, T., Farmer, R., Gabernet, G., Lucas, C.L., et al. (2025). A single-cell atlas of circulating immune cells over the first 2 months of age in extremely premature infants. Sci. Transl. Med. 17, eadr0942. 10.1126/scitranslmed.adr0942.

10. Fu, Y., Wen, Z., and Fan, J. (2025). Interaction of low-density neutrophils with other immune cells in the mechanism of inflammation. Mol. Med. 31, 133. 10.1186/s10020-025-01187-5.

11. Köstlin, N., Vogelmann, M., Spring, B., Schwarz, J., Feucht, J., Härtel, C., Orlikowsky, T.W., Poets, C.F., and Gille, C. (2017). Granulocytic myeloid-derived suppressor cells from human cord blood modulate T-helper cell response towards an anti-inflammatory phenotype. Immunology 152, 89–101. 10.1111/imm.12751.

12. Gervassi, A., Lejarcegui, N., Dross, S., Jacobson, A., Itaya, G., Kidzeru, E., Gantt, S., Jaspan, H., and Horton, H. (2014). Myeloid Derived Suppressor Cells Are Present at High Frequency in Neonates and Suppress In Vitro T Cell Responses. PLOS ONE 9, e107816. 10.1371/journal.pone.0107816.

13. He, Y.-M., Li, X., Perego, M., Nefedova, Y., Kossenkov, A.V., Jensen, E.A., Kagan, V., Liu, Y.-F., Fu, S.-Y., Ye, Q.-J., et al. (2018). Transitory presence of myeloid-derived suppressor cells in neonates is critical for control of inflammation. Nat. Med. 24, 224–231. 10.1038/nm.4467.

14. Ulas, T., Pirr, S., Fehlhaber, B., Bickes, M.S., Loof, T.G., Vogl, T., Mellinger, L., Heinemann, A.S., Burgmann, J., Schöning, J., et al. (2017). S100-alarmin-induced innate immune programming protects newborn infants from sepsis. Nat. Immunol. 18, 622–632. 10.1038/ni.3745.

15. Lee, A.H., Shannon, C.P., Amenyogbe, N., Bennike, T.B., Diray-Arce, J., Idoko, O.T., Gill, E.E., Ben-Othman, R., Pomat, W.S., van Haren, S.D., et al. (2019). Dynamic molecular changes during the first week of human life follow a robust developmental trajectory. Nat. Commun. 10, 1092. 10.1038/s41467-019-08794-x.

16. Nehar-Belaid, D., Thibodeau, A., Eroglu, A., Marches, R., Eryilmaz, G., Unutmaz, D., Verschoor, C.P., Gu, J., Balaji, U., Mejías, A., et al. (2025). Single-cell map of the healthy human immune system across the lifespan reveals unique infant immune signatures. BioRxiv Prepr. Serv. Biol., 2025.07.28.667181. 10.1101/2025.07.28.667181.

17. Wang, G., Miyahara, Y., Guo, Z., Khattar, M., Stepkowski, S.M., and Chen, W. (2010). “Default” generation of neonatal regulatory T cells. J. Immunol. Baltim. Md 1950 185, 71–78. 10.4049/jimmunol.0903806.

18. Bunis, D.G., Bronevetsky, Y., Krow-Lucal, E., Bhakta, N.R., Kim, C.C., Nerella, S., Jones, N., Mendoza, V.F., Bryson, Y.J., Gern, J.E., et al. (2021). Single-Cell Mapping of Progressive Fetal-to-Adult Transition in Human Naive T Cells. Cell Rep. 34, 108573. 10.1016/j.celrep.2020.108573.

19. Hebel, K., Weinert, S., Kuropka, B., Knolle, J., Kosak, B., Jorch, G., Arens, C., Krause, E., Braun-Dullaeus, R.C., and Brunner-Weinzierl, M.C. (2014). CD4+ T cells from human neonates and infants are poised spontaneously to run a nonclassical IL-4 program. J. Immunol. Baltim. Md 1950 192, 5160–5170. 10.4049/jimmunol.1302539.

20. Wu, C.Y., Demeure, C., Kiniwa, M., Gately, M., and Delespesse, G. (1993). IL-12 induces the production of IFN-gamma by neonatal human CD4 T cells. J. Immunol. Baltim. Md 1950 151, 1938–1949.

21. Watson, N.B., Patel, R.K., Kean, C., Veazey, J., Oyesola, O.O., Laniewski, N., Grenier, J.K., Wang, J., Tabilas, C., Yee Mon, K.J., et al. (2024). The gene regulatory basis of bystander activation in CD8+ T cells. Sci. Immunol. 9, eadf8776. 10.1126/sciimmunol.adf8776.

22. Ehlers, G., Tödtmann, A.M., Holsten, L., Willers, M., Heckmann, J., Schöning, J., Richter, M., Heinemann, A.S., Pirr, S., Heinz, A., et al. (2025). Oxidative phosphorylation is a key feature of neonatal monocyte immunometabolism promoting myeloid differentiation after birth. Nat. Commun. 16, 2239. 10.1038/s41467-025-57357-w.

23. Büttner, M., Ostner, J., Müller, C.L., Theis, F.J., and Schubert, B. (2021). scCODA is a Bayesian model for compositional single-cell data analysis. Nat. Commun. 12, 6876. 10.1038/s41467-021-27150-6.

24. St Laurent, C.D., Jame-Chenarboo, Z., Beck, A.E., Stubblefield, S., Duan, S., Sigal, D., and Macauley, M.S. (2025). CD16 and Siglec expression refine the phenotypic heterogeneity of steady-state myeloid-derived suppressor cells. Front. Oncol. 15, 1570121. 10.3389/fonc.2025.1570121.

25. Sciumè, G., De Angelis, G., Benigni, G., Ponzetta, A., Morrone, S., Santoni, A., and Bernardini, G. (2011). CX3CR1 expression defines 2 KLRG1+ mouse NK-cell subsets with distinct functional properties and positioning in the bone marrow. Blood 117, 4467–4475. 10.1182/blood-2010-07-297101.

26. Kim, D.O., Byun, J.-E., Kim, W.S., Kim, M.J., Choi, J.H., Kim, H., Choi, E., Kim, T.-D., Yoon, S.R., Noh, J.-Y., et al. (2020). TXNIP Regulates Natural Killer Cell-Mediated Innate Immunity by Inhibiting IFN-γ Production during Bacterial Infection. Int. J. Mol. Sci. 21, 9499. 10.3390/ijms21249499.

27. Cubitt, C.C., Wong, P., Dorando, H.K., Foltz, J.A., Tran, J., Marsala, L., Marin, N.D., Foster, M., Schappe, T., Fatima, H., et al. (2024). Induced CD8α identifies human NK cells with enhanced proliferative fitness and modulates NK cell activation. J. Clin. Invest. 134, e173602. 10.1172/JCI173602.

28. McKinney, E.F., Cuthbertson, I., Harris, K.M., Smilek, D.E., Connor, C., Manferrari, G., Carr, E.J., Zamvil, S.S., and Smith, K.G.C. (2021). A CD8+ NK cell transcriptomic signature associated with clinical outcome in relapsing remitting multiple sclerosis. Nat. Commun. 12, 635. 10.1038/s41467-020-20594-2.

29. Brodsky, N.N., Chakder, M., Kumar, D.B.U., Barmada, A., Wang, J., Gu, W., Olaloye, O., Ramaswamy, A., Wats, A., Konnikova, L., et al. (2025). Early-life human CD8+ T cells exhibit rapid, short-lived effector responses and a unique transcription factor landscape. Proc. Natl. Acad. Sci. U. S. A. 122, e2421106122. 10.1073/pnas.2421106122.

30. Cheng, T.-L., Lai, C.-H., Shieh, S.-J., Jou, Y.-B., Yeh, J.-L., Yang, A.-L., Wang, Y.-H., Wang, C.-Z., Chen, C.-H., Shi, G.-Y., et al. (2016). Myeloid thrombomodulin lectin-like domain inhibits osteoclastogenesis and inflammatory bone loss. Sci. Rep. 6, 28340. 10.1038/srep28340.

31. Janssen, L.L.G., van Leeuwen-Kerkhoff, N., Westers, T.M., de Gruijl, T.D., and van de Loosdrecht, A.A. (2024). The immunoregulatory role of monocytes and thrombomodulin in myelodysplastic neoplasms. Front. Oncol. 14. 10.3389/fonc.2024.1414102.

32. Riffelmacher, T., Giles, D.A., Zahner, S., Dicker, M., Andreyev, A.Y., McArdle, S., Perez-Jeldres, T., van der Gracht, E., Murray, M.P., Hartmann, N., et al. (2021). Metabolic activation and colitis pathogenesis is prevented by lymphotoxin β receptor expression in neutrophils. Mucosal Immunol. 14, 679–690. 10.1038/s41385-021-00378-7.

33. Ip, W.K.E., Hoshi, N., Shouval, D.S., Snapper, S., and Medzhitov, R. (2017). Anti-inflammatory effect of IL-10 mediated by metabolic reprogramming of macrophages. Science 356, 513–519. 10.1126/science.aal3535.

34. Jiao, Y., and Xiang, Y. (2025). A review of the participation of DDIT4 in the tumor immune microenvironment through inhibiting PI3K-Akt/mTOR pathway. Front. Oncol. 15, 1595463. 10.3389/fonc.2025.1595463.

35. Cuenca, A.G., Wynn, J.L., Kelly-Scumpia, K.M., Scumpia, P.O., Vila, L., Delano, M.J., Mathews, C.E., Wallet, S.M., Reeves, W.H., Behrns, K.E., et al. (2011). Critical Role for CXC Ligand 10/CXC Receptor 3 Signaling in the Murine Neonatal Response to Sepsis. Infect. Immun. 79, 2746–2754. 10.1128/iai.01291-10.

36. Peperzak, V., Veraar, E.A.M., Xiao, Y., Babala, N., Thiadens, K., Brugmans, M., and Borst, J. (2013). CD8+ T cells produce the chemokine CXCL10 in response to CD27/CD70 costimulation to promote generation of the CD8+ effector T cell pool. J. Immunol. Baltim. Md 1950 191, 3025–3036. 10.4049/jimmunol.1202222.

37. Troulé, K., Petryszak, R., Cakir, B., Cranley, J., Harasty, A., Prete, M., Tuong, Z.K., Teichmann, S.A., Garcia-Alonso, L., and Vento-Tormo, R. (2025). CellPhoneDB v5: inferring cell–cell communication from single-cell multiomics data. Nat. Protoc., 1–29. 10.1038/s41596-024-01137-1.

38. Liu, S., Zhou, Y., Li, G., Zhu, B., Wu, F., Zhou, J., Chen, X., Qin, B., Gao, Y., Wang, F., et al. (2025). PLAUR+ Neutrophils Drive Anti-PD-1 Therapy Resistance in Patients with Hepatocellular Carcinoma by Shaping an Immunosuppressive Microenvironment. Adv. Sci. Weinh. Baden-Wurtt. Ger. 12, e07167. 10.1002/advs.202507167.

39. Lugassy, J., Abdala-Saleh, N., Jarrous, G., Turky, A., Saidemberg, D., Ridner-Bahar, G., Berger, N., Bar-On, D., Taura, T., Wilson, D., et al. Development of DPP-4-resistant CXCL9-Fc and CXCL10-Fc chemokines for effective cancer immunotherapy. Proc. Natl. Acad. Sci. U. S. A. 122, e2501791122. 10.1073/pnas.2501791122.

40. Fallone, L., Pouxvielh, K., Arbez, L., Rousseaux, N., Picq, L., Drouillard, A., Mathieu, A.-L., Nombel, A., Benezech, S., Bourdonnay, E., et al. (2025). Interleukins 15 and 18 synergistically prime the antitumor function of natural killer cells through noncanonical activation of mTORC1. Sci. Signal. 18, eadq8778. 10.1126/scisignal.adq8778.

41. Berjis, A., Muthumani, D., Aguilar, O.A., Pomp, O., Johnson, O., Finck, A.V., Engel, N.W., Chen, L., Plachta, N., Scholler, J., et al. (2024). Pretreatment with IL-15 and IL-18 rescues natural killer cells from granzyme B-mediated apoptosis after cryopreservation. Nat. Commun. 15, 3937. 10.1038/s41467-024-47574-0.

42. Levanon, D., Negreanu, V., Lotem, J., Bone, K.R., Brenner, O., Leshkowitz, D., and Groner, Y. (2014). Transcription factor Runx3 regulates interleukin-15-dependent natural killer cell activation. Mol. Cell. Biol. 34, 1158–1169. 10.1128/MCB.01202-13.

43. Rea, A., Santana-Hernández, S., Villanueva, J., Sanvicente-García, M., Cabo, M., Suarez-Olmos, J., Quimis, F., Qin, M., Llorens, E., Blasco-Benito, S., et al. (2025). Enhancing human NK cell antitumor function by knocking out SMAD4 to counteract TGFβ and activin A suppression. Nat. Immunol. 26, 582–594. 10.1038/s41590-025-02103-z.

44. Viel, S., Marçais, A., Guimaraes, F.S.-F., Loftus, R., Rabilloud, J., Grau, M., Degouve, S., Djebali, S., Sanlaville, A., Charrier, E., et al. (2016). TGF-β inhibits the activation and functions of NK cells by repressing the mTOR pathway. Sci. Signal. 9, ra19–ra19. 10.1126/scisignal.aad1884.

45. Wehner, R., Löbel, B., Bornhäuser, M., Schäkel, K., Cartellieri, M., Bachmann, M., Rieber, E.P., and Schmitz, M. (2009). Reciprocal activating interaction between 6-sulfo LacNAc+ dendritic cells and NK cells. Int. J. Cancer 124, 358–366. 10.1002/ijc.23962.

46. Wang, J., Zhao, X., and Wan, Y.Y. (2023). Intricacies of TGF-β signaling in Treg and Th17 cell biology. Cell. Mol. Immunol. 20, 1002–1022. 10.1038/s41423-023-01036-7.

47. Fantini, M.C., Becker, C., Monteleone, G., Pallone, F., Galle, P.R., and Neurath, M.F. (2004). Cutting edge: TGF-beta induces a regulatory phenotype in CD4+CD25- T cells through Foxp3 induction and down-regulation of Smad7. J. Immunol. Baltim. Md 1950 172, 5149–5153. 10.4049/jimmunol.172.9.5149.

48. Laouar, Y., Sutterwala, F.S., Gorelik, L., and Flavell, R.A. (2005). Transforming growth factor-β controls T helper type 1 cell development through regulation of natural killer cell interferon-γ. Nat. Immunol. 6, 600–607. 10.1038/ni1197.

49. Oesinghaus, L., Becker, S., Vornholz, L., Papalexi, E., Pangallo, J., Moinfar, A.A., Liu, J., Fleur, A.L., Shulman, M., Marrujo, S., et al. (2025). A single-cell cytokine dictionary of human peripheral blood. bioRxiv, 2025.12.12.693897. 10.64898/2025.12.12.693897.

50. Schulz, E.G., Mariani, L., Radbruch, A., and Höfer, T. (2009). Sequential Polarization and Imprinting of Type 1 T Helper Lymphocytes by Interferon-γ and Interleukin-12. Immunity 30, 673–683. 10.1016/j.immuni.2009.03.013.

51. Feili-Hariri, M., Falkner, D.H., and Morel, P.A. (2005). Polarization of naive T cells into Th1 or Th2 by distinct cytokine-driven murine dendritic cell populations: implications for immunotherapy. J. Leukoc. Biol. 78, 656–664. 10.1189/jlb.1104631.

52. Ben-Sasson, S.Z., Hu-Li, J., Quiel, J., Cauchetaux, S., Ratner, M., Shapira, I., Dinarello, C.A., and Paul, W.E. (2009). IL-1 acts directly on CD4 T cells to enhance their antigen-driven expansion and differentiation. Proc. Natl. Acad. Sci. 106, 7119–7124. 10.1073/pnas.0902745106.

53. Griffith, J.W., Faustino, L.D., Cottrell, V.I., Nepal, K., Hariri, L.P., Chiu, R.S.-Y., Jones, M.C., Julé, A., Gabay, C., and Luster, A.D. (2023). Regulatory T cell-derived IL-1Ra suppresses the innate response to respiratory viral infection. Nat. Immunol. 24, 2091–2107. 10.1038/s41590-023-01655-2.

54. Ermann, J., and Fathman, C.G. (2003). Costimulatory signals controlling regulatory T cells. Proc. Natl. Acad. Sci. 100, 15292–15293. 10.1073/pnas.0307001100.

55. Kim, J.I., Sonawane, S.B., Lee, M.K., Lee, S.-H., Duff, P.E., Moore, D.J., O’Connor, M.R., Lian, M.-M., Deng, S., Choi, Y., et al. (2010). Blockade of GITR–GITRL interaction maintains Treg function to prolong allograft survival. Eur. J. Immunol. 40, 1369–1374. 10.1002/eji.200940046.

56. Lin, L., Spoor, M.S., Gerth, A.J., Brody, S.L., and Peng, S.L. (2004). Modulation of Th1 Activation and Inflammation by the NF-κB Repressor Foxj1. Science 303, 1017–1020. 10.1126/science.1093889.

57. Liu, X., Wang, Y., Lu, H., Li, J., Yan, X., Xiao, M., Hao, J., Alekseev, A., Khong, H., Chen, T., et al. (2019). Genome-wide analysis identifies NR4A1 as a key mediator of T cell dysfunction. Nature 567, 525–529. 10.1038/s41586-019-0979-8.

58. Lohoff, M., Giaisi, M., Köhler, R., Casper, B., Krammer, P.H., and Li-Weber, M. (2010). Early Growth Response Protein-1 (Egr-1) Is Preferentially Expressed in T Helper Type 2 (Th2) Cells and Is Involved in Acute Transcription of the Th2 Cytokine Interleukin-4. J. Biol. Chem. 285, 1643–1652. 10.1074/jbc.M109.011585.

59. Zhao, Y., Xiao, C., Li, S., Huang, A., Li, H., Dong, J., Qu, Q., Liu, X., Gao, B., and Shao, N. (2025). CD71-Mediated Effects of Soluble Vasorin on Tumor Progression, Angiogenesis and Immunosuppression. Int. J. Mol. Sci. 26, 4913. 10.3390/ijms26104913.

60. Tang, X., Sui, X., Weng, L., and Liu, Y. (2021). SNAIL1: Linking Tumor Metastasis to Immune Evasion. Front. Immunol. 12. 10.3389/fimmu.2021.724200.

61. Joller, N., Lozano, E., Burkett, P.R., Patel, B., Xiao, S., Zhu, C., Xia, J., Tan, T.G., Sefik, E., Yajnik, V., et al. (2014). Treg cells expressing the co-inhibitory molecule TIGIT selectively inhibit pro-inflammatory Th1 and Th17 cell responses. Immunity 40, 569–581. 10.1016/j.immuni.2014.02.012.

62. Yu, X., Harden, K., Gonzalez, L.C., Francesco, M., Chiang, E., Irving, B., Tom, I., Ivelja, S., Refino, C.J., Clark, H., et al. (2009). The surface protein TIGIT suppresses T cell activation by promoting the generation of mature immunoregulatory dendritic cells. Nat. Immunol. 10, 48–57. 10.1038/ni.1674.

63. Muri, J., Thut, H., and Kopf, M. (2021). The thioredoxin-1 inhibitor Txnip restrains effector T-cell and germinal center B-cell expansion. Eur. J. Immunol. 51, 115–124. 10.1002/eji.202048851.

64. Willcox, A.C., Gobillot, T.A., Kikawa, C., Baumgarten, N.E., Stoddard, C.I., Sung, K., Bhattacharya, T., Freeman, T.S., Marceau, J., Humes, D., et al. (2025). Identification of AMOTL2 as an antiviral factor that enhances the human type I interferon response against Zika virus. Proc. Natl. Acad. Sci. 122, e2507955122. 10.1073/pnas.2507955122.

65. Basole, C.P., Nguyen, R.K., Lamothe, K., Vang, A., Clark, R., Baillie, G.S., Epstein, P.M., and Brocke, S. (2017). PDE8 controls CD4+ T cell motility through the PDE8A-Raf-1 kinase signaling complex. Cell. Signal. 40, 62–72. 10.1016/j.cellsig.2017.08.007.

66. Epstein, P.M., Basole, C., and Brocke, S. (2021). The Role of PDE8 in T Cell Recruitment and Function in Inflammation. Front. Cell Dev. Biol. 9, 636778. 10.3389/fcell.2021.636778.

67. Croft, M., So, T., Duan, W., and Soroosh, P. (2009). The Significance of OX40 and OX40L to T cell Biology and Immune Disease. Immunol. Rev. 229, 173–191. 10.1111/j.1600-065X.2009.00766.x.

68. Yu, F., Sharma, S., Jankovic, D., Gurram, R.K., Su, P., Hu, G., Li, R., Rieder, S., Zhao, K., Sun, B., et al. (2018). The transcription factor Bhlhe40 is a switch of inflammatory versus antiinflammatory Th1 cell fate determination. J. Exp. Med. 215, 1813–1821. 10.1084/jem.20170155.

69. Smith, G.A., Taunton, J., and Weiss, A. (2017). IL-2Rβ Abundance Differentially Tunes IL-2 Signaling Dynamics in CD4+ and CD8+ T Cells. Sci. Signal. 10, eaan4931. 10.1126/scisignal.aan4931.

70. Miyara, M., Yoshioka, Y., Kitoh, A., Shima, T., Wing, K., Niwa, A., Parizot, C., Taflin, C., Heike, T., Valeyre, D., et al. (2009). Functional delineation and differentiation dynamics of human CD4+ T cells expressing the FoxP3 transcription factor. Immunity 30, 899–911. 10.1016/j.immuni.2009.03.019.

71. Herzenberg, L.A., and Herzenberg, L.A. (1989). Toward a layered immune system. Cell 59, 953–954. 10.1016/0092-8674(89)90748-4.

72. Yoshida, M., Worlock, K.B., Huang, N., Lindeboom, R.G.H., Butler, C.R., Kumasaka, N., Dominguez Conde, C., Mamanova, L., Bolt, L., Richardson, L., et al. (2022). Local and systemic responses to SARS-CoV-2 infection in children and adults. Nature 602, 321–327. 10.1038/s41586-021-04345-x.

73. Li, Y., Kong, W., Yang, W., Patel, R.M., Casey, E.B., Okeyo-Owuor, T., White, J.M., Porter, S.N., Morris, S.A., and Magee, J.A. (2020). Single-Cell Analysis of Neonatal HSC Ontogeny Reveals Gradual and Uncoordinated Transcriptional Reprogramming that Begins before Birth. Cell Stem Cell 27, 732–747.e7. 10.1016/j.stem.2020.08.001.

74. Liu, Y., Perego, M., Xiao, Q., He, Y., Fu, S., He, J., Liu, W., Li, X., Tang, Y., Li, X., et al. Lactoferrin-induced myeloid-derived suppressor cell therapy attenuates pathologic inflammatory conditions in newborn mice. J. Clin. Invest. 129, 4261–4275. 10.1172/JCI128164.

75. Schwarz, J., Scheckenbach, V., Kugel, H., Spring, B., Pagel, J., Härtel, C., Pauluschke-Fröhlich, J., Peter, A., Poets, C.F., Gille, C., et al. (2018). Granulocytic myeloid-derived suppressor cells (GR-MDSC) accumulate in cord blood of preterm infants and remain elevated during the neonatal period. Clin. Exp. Immunol. 191, 328–337. 10.1111/cei.13059.

76. Alicea-Torres, K., Sanseviero, E., Gui, J., Chen, J., Veglia, F., Yu, Q., Donthireddy, L., Kossenkov, A., Lin, C., Fu, S., et al. (2021). Immune suppressive activity of myeloid-derived suppressor cells in cancer requires inactivation of the type I interferon pathway. Nat. Commun. 12, 1717. 10.1038/s41467-021-22033-2.

77. Sun, Y., Han, X., Shang, C., Wang, Y., Xu, B., Jiang, S., Mo, Y., Wang, D., Ke, Y., and Zeng, X. (2022). The downregulation of type I IFN signaling in G-MDSCs under tumor conditions promotes their development towards an immunosuppressive phenotype. Cell Death Dis. 13, 36. 10.1038/s41419-021-04487-w.

78. Collins, A., Swann, J.W., Proven, M.A., Patel, C.M., Mitchell, C.A., Kasbekar, M., Dellorusso, P.V., and Passegué, E. (2024). Maternal inflammation regulates fetal emergency myelopoiesis. Cell 187, 1402–1421.e21. 10.1016/j.cell.2024.02.002.

79. Klimchenko, O., Di Stefano, A., Geoerger, B., Hamidi, S., Opolon, P., Robert, T., Routhier, M., El-Benna, J., Delezoide, A.-L., Boukour, S., et al. (2011). Monocytic cells derived from human embryonic stem cells and fetal liver share common differentiation pathways and homeostatic functions. Blood 117, 3065–3075. 10.1182/blood-2010-07-295246.

80. Mold, J.E., Venkatasubrahmanyam, S., Burt, T.D., Michaëlsson, J., Rivera, J.M., Galkina, S.A., Weinberg, K., Stoddart, C.A., and McCune, J.M. (2010). Fetal and adult hematopoietic stem cells give rise to distinct T cell lineages in humans. Science 330, 1695–1699. 10.1126/science.1196509.

81. Neavin, D., Senabouth, A., Arora, H., Lee, J.T.H., Ripoll-Cladellas, A., sc-eQTLGen Consortium, Franke, L., Prabhakar, S., Ye, C.J., McCarthy, D.J., et al. (2024). Demuxafy: improvement in droplet assignment by integrating multiple single-cell demultiplexing and doublet detection methods. Genome Biol. 25, 94. 10.1186/s13059-024-03224-8.

82. Germain, P.-L., Lun, A., Garcia Meixide, C., Macnair, W., and Robinson, M.D. (2021). Doublet identification in single-cell sequencing data using scDblFinder. F1000Research 10, 979. 10.12688/f1000research.73600.2.

83. Gayoso, A., Steier, Z., Lopez, R., Regier, J., Nazor, K.L., Streets, A., and Yosef, N. (2021). Joint probabilistic modeling of single-cell multi-omic data with totalVI. Nat. Methods 18, 272–282. 10.1038/s41592-020-01050-x.

84. Wolf, F.A., Angerer, P., and Theis, F.J. (2018). SCANPY: large-scale single-cell gene expression data analysis. Genome Biol. 19, 15. 10.1186/s13059-017-1382-0.

85. Hao, Y., Stuart, T., Kowalski, M.H., Choudhary, S., Hoffman, P., Hartman, A., Srivastava, A., Molla, G., Madad, S., Fernandez-Granda, C., et al. (2024). Dictionary learning for integrative, multimodal and scalable single-cell analysis. Nat. Biotechnol. 42, 293–304. 10.1038/s41587-023-01767-y.

86. Hua, Y., Weng, L., Zhao, F., and Rambow, F. (2025). SeuratExtend: streamlining single-cell RNA-seq analysis through an integrated and intuitive framework. GigaScience 14, giaf076. 10.1093/gigascience/giaf076.

87. Miao, B.-B., Dong, W., Han, Z.-F., Luo, X., Ke, C.-H., and You, W.-W. (2023). TOmicsVis: An all-in-one transcriptomic analysis and visualization R package with Shinyapp interface. iMeta 2, e137. 10.1002/imt2.137.

88. Chen, E.Y., Tan, C.M., Kou, Y., Duan, Q., Wang, Z., Meirelles, G.V., Clark, N.R., and Ma’ayan, A. (2013). Enrichr: interactive and collaborative HTML5 gene list enrichment analysis tool. BMC Bioinformatics 14, 128. 10.1186/1471-2105-14-128.

89. Yu, G., Wang, L.-G., Han, Y., and He, Q.-Y. (2012). clusterProfiler: an R Package for Comparing Biological Themes Among Gene Clusters. OMICS J. Integr. Biol. 16, 284–287. 10.1089/omi.2011.0118.

