## Supplementary material for "Human neonatal CITE-seq atlas identifies an immune transition at 32 weeks’ gestation from CD15⁺ myeloid-dominated to interferon-primed immunity": Document S1

### MATERIALS and METHODS

#### *Cell isolation and cryopreservation*

Cord blood mononuclear cells (CBMCs) and peripheral blood mononuclear cells (PBMCs) were isolated by density gradient centrifugation using Biocoll (Bio&SELL). Whole blood was layered onto Biocoll at a 1:2 ratio (blood:Biocoll) and centrifuged at 1600 rpm for 20 min at room temperature without a break. Cells were resuspended in PBS + 2% fetal calf serum (FCS) at a volume corresponding to the original blood volume. Residual erythrocytes were depleted using EasySep™ RBC Depletion Reagent (STEMCELL Technologies) according to the manufacturer's instructions, with minor adjustments in bead volume. To minimize monocytes loss during magnetic separation, EDTA (Invitrogen by ThermoFisher SCIENTIFIC) was added to a final concentration of 6mM prior to labeling. Beads were vortexed for 30 s and added proportionally to sample volume (25 µL/mL), followed by 5 min incubation at room temperature in an EasyEights™ EasySep™ Magnet (STEMCELL Technologies). The supernatant containing unlabeled cells was retained, washed once with PBS + 2% FCS, and centrifuged at 400 x g for 10 min. Cell numbers and viability were determined using a Countess automated cell counter (ThermoFisher SCIENTIFIC) after 1:1 dilution with Trypan Blue. For cryopreservation, cells were resuspended in chilled resuspension medium, mixed 1:1 with chilled 2xfreezing medium to obtain a final composition of 40% FCS and 15% DMSO, aliquoted at 1 - 10 x $10^6$ cells/mL, placed in a pre-cooled freezing container at -80°C for 24 h, and subsequently transferred to liquid nitrogen until further processing.

SUPPLEMENTARY FIGURES

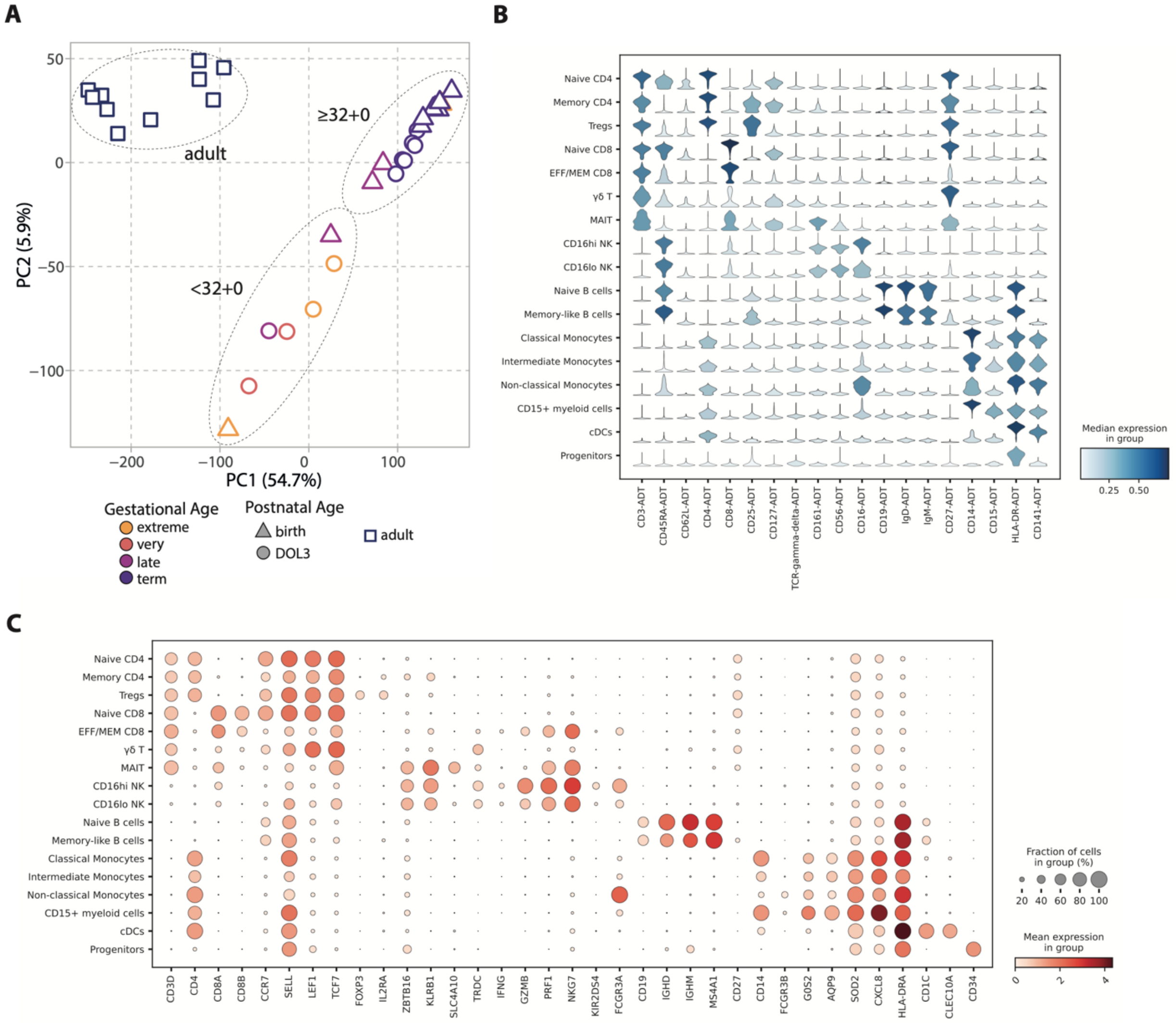


**Figure S1.**
(A) PCA of normalized RNA expression, aggregated by donor. (B) Violin plot showing median normalized ADT marker expression in each cell subpopulation. (C) Dot plot showing mean normalized RNA marker expression in each cell subset. ADT: Antibody-derived tag.

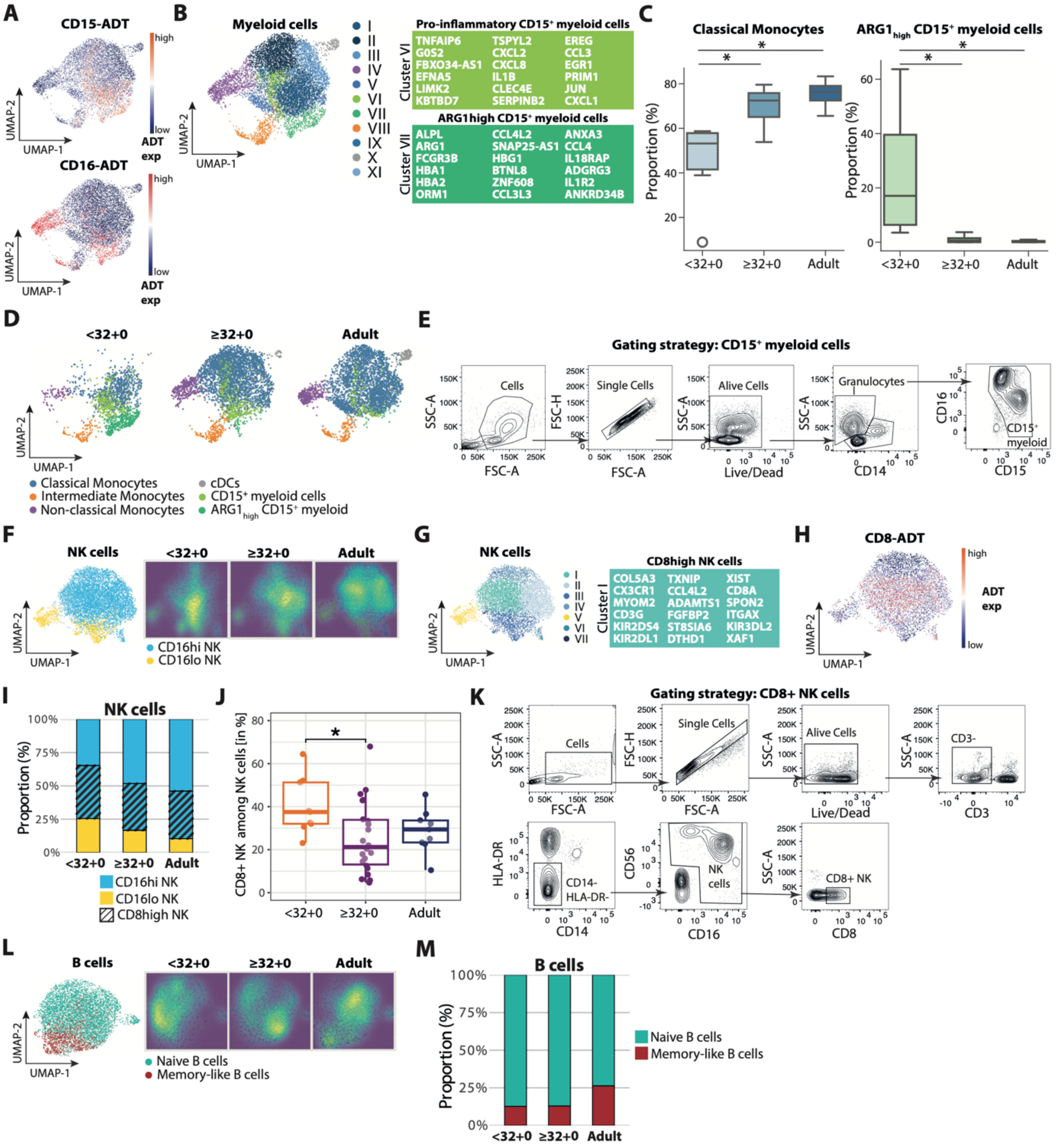


**Figure S2.**
(A) CD15-ADT (top) and CD16-ADT (bottom) feature plots of myeloid cells at baseline, derived from neonatal birth and DOL3, as well as adult samples. (B) UMAP of myeloid cells at baseline, derived from neonatal birth and DOL3, as well as adult samples with two unsupervised CD15$^+$ myeloid cell subclusters; top markers for each cluster highlighted. (C) Boxplots show the proportion of classical monocytes (left) and ARG1$_{high}$CD15$^+$ myeloid cells (right) among myeloid cells per sample. ARG1$_{high}$CD15$^+$ myeloid cells were credibly increased, and classical monocytes were credibly decreased in <32+0 compared to ≥32+0-wk and adult samples, as determined by Bayesian compositional analysis using scCODA[23] (intermediate monocytes as reference). Asterisks indicate cell types identified as credibly altered by scCODA and do not represent p values. (D) UMAP plots of myeloid cells split by age group. (E) Flow cytometry gating strategy for CD15$^+$ myeloid cells. (F) Left: UMAP plot of NK cells, derived from neonatal birth and DOL3, as well as adult samples. Right: Normalized density UMAPs of NK cells split by age group. (G) UMAP of NK cells at baseline, derived from neonatal birth and DOL3, as well as adult samples with unsupervised subclusters; top markers for cluster I highlighted. (H) CD8-ADT feature plot of NK cells at baseline, derived from neonatal birth and DOL3, as well as adult samples. (I) Stacked bar graph representing the relative proportion of cells per cell type and group among NK cells. (J) CD8+ NK cell frequency among NK cells grouped by age group as assessed by flow cytometry (<32-wk, n=9; ≥32+0-wk, n=22; adults, n=9; one-way ANOVA; p: * < 0.05). (K) Flow cytometry gating strategy for CD8+ NK cells. (L) UMAP plot of B cells at baseline across age groups, as in (F) for NK cells. (M) Relative proportions of B cell subsets across age groups, as in (I) for NK cells.

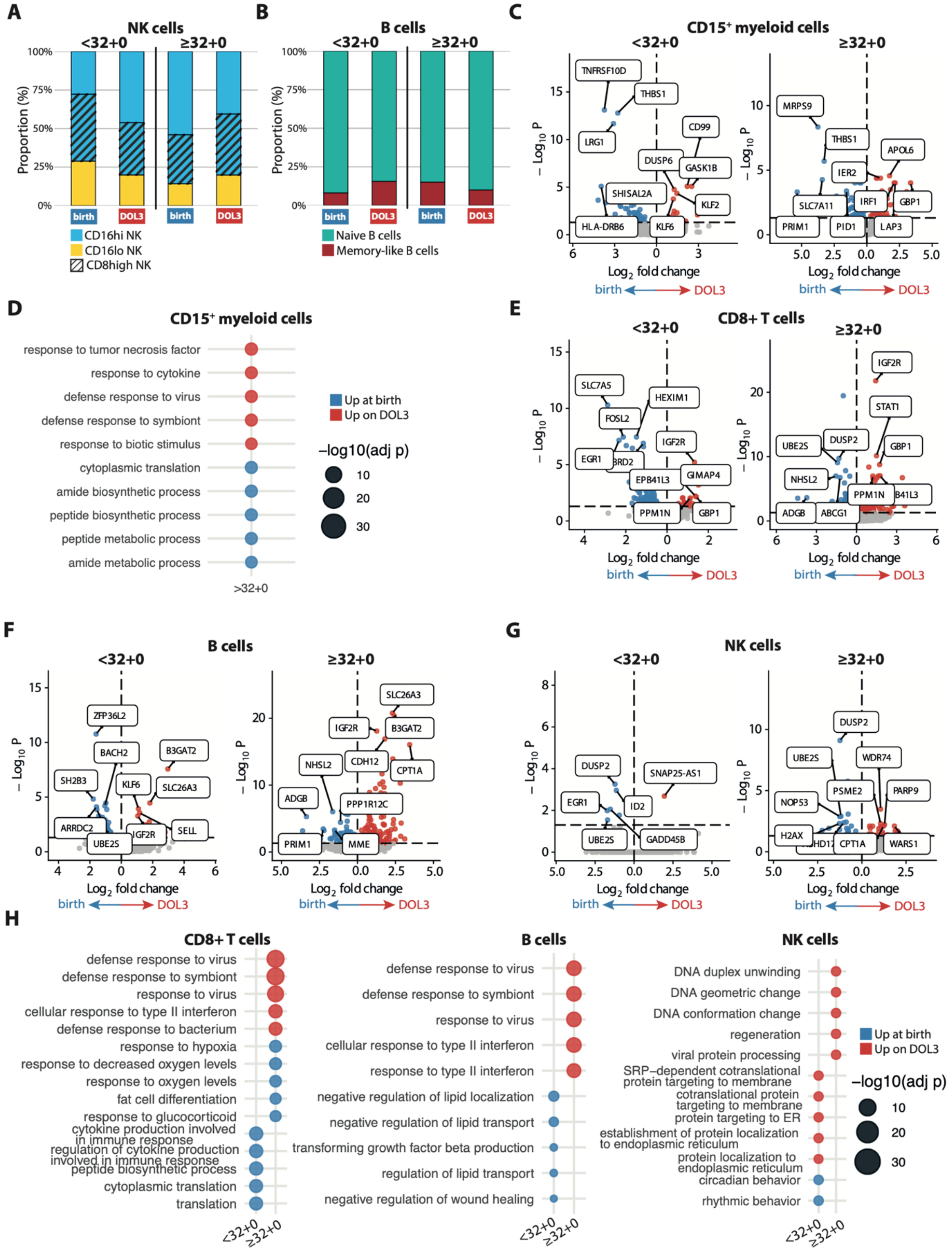


**Figure S3.**
(A-B) Stacked bar graph representing the relative proportion of cells per cell type and group among NK cells (A) and B cells (B). (C) Volcano plots of baseline gene expression in neonatal CD15$^{+}$ myeloid cells (birth vs. DOL3), stratified by age group (adj. $p < 0.05$). Top DEGs (|log2FC| > 1) were labeled based on ranked adjusted p values. (D) Dot plot of enriched GO Biological Process terms in CD15$^{+}$ myeloid cells for neonates >32+0 weeks, based on genes upregulated at birth or on DOL3 (adj. $p < 0.05$), respectively, with the top 5 terms per condition ranked by adjusted p value (adj. $p < 0.05$). No significant GO term enrichment was detected for the <32+0 weeks groups. (E-G) Volcano plots of baseline gene expression in neonatal CD8+ T cells (E), B cells (F), and NK cells (G) (birth vs. DOL3), analogous to (C) for CD15$^{+}$ myeloid cells (H) Dot plot of GO Biological Process enrichment in CD8+ T cells, B cells, and NK cells for neonates ≥32+0 and <32+0 weeks, analogous to (D) for CD15$^{+}$ myeloid cells.

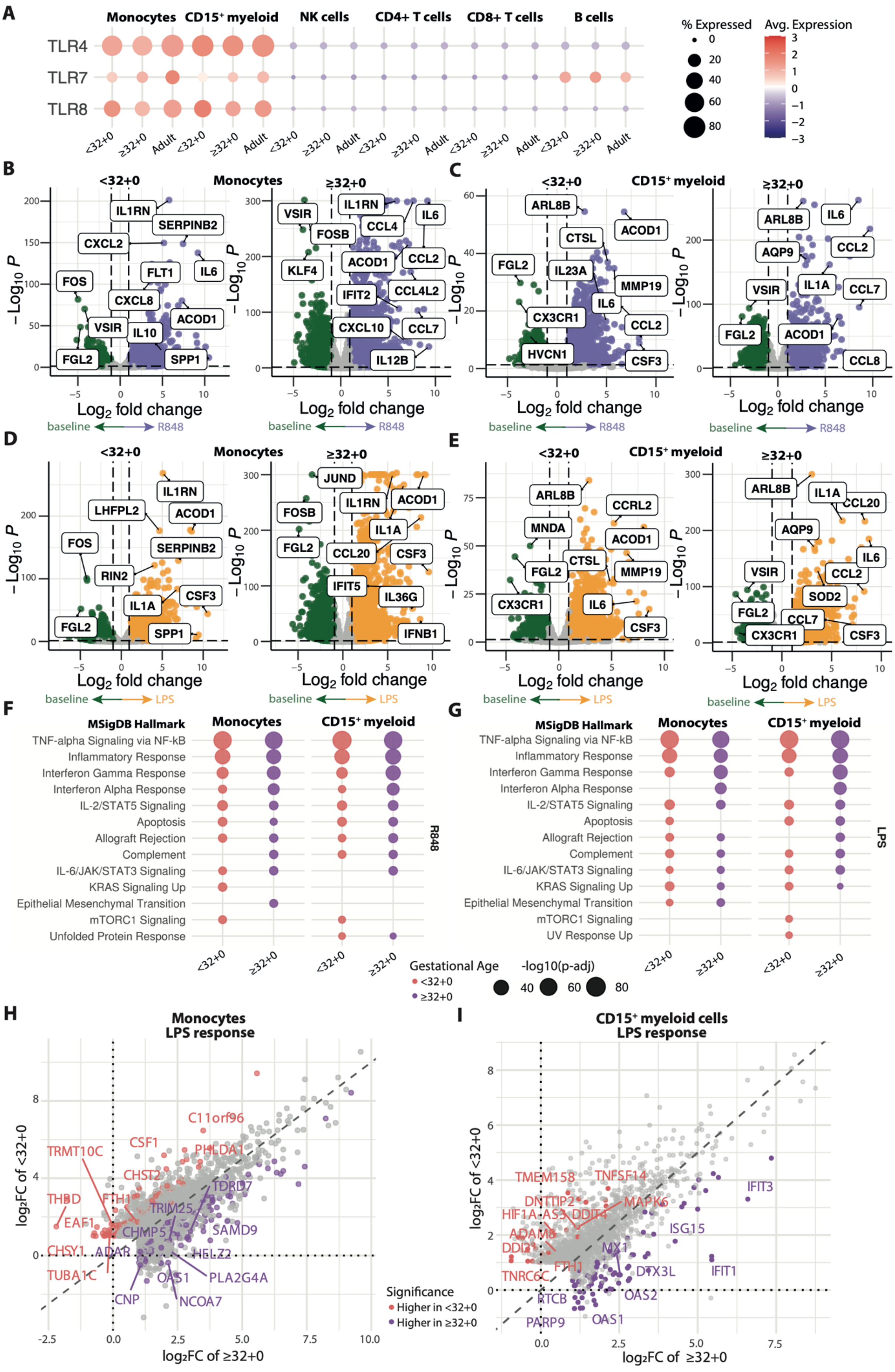
A
Monocytes
CD15⁺ myeloid
NK cells
CD4+ T cells
CD8+ T cells
B cells
TLR4
TLR7
TLR8
<32+0
≥32+0
Adult
% Expressed
0
20
40
60
80
Avg. Expression
3
2
1
0
−1
−2
−3
B
Monocytes
<32+0
≥32+0
IL1RN
SERPINB2
CXCL2
FLT1
IL6
FOS
CXCL8
ACOD1
VSIR
IL10
FGL2
SPP1
VSIR
FOSB
CCL4
KLF4
ACOD1
CCL2
IFIT2
CCL4L2
CXCL10
CCL7
IL12B
−Log10 P
Log2 fold change
baseline
R848
C
CD15⁺ myeloid
ARL8B
ACOD1
CTSL
FGL2
IL23A
MMP19
IL6
CX3CR1
CCL2
HVCN1
CSF3
ARL8B
AQP9
CCL2
IL1A
CCL7
FGL2
ACOD1
CCL8
D
Monocytes
IL1RN
LHFPL2
ACOD1
FOS
SERPINB2
RIN2
IL1A
CSF3
FGL2
SPP1
JUND
FOSB
IL1RN
ACOD1
IL1A
FGL2
CCL20
CSF3
IFIT5
IL36G
IFNB1
LPS
E
CD15⁺ myeloid
ARL8B
CCRL2
MNDA
ACOD1
CTSL
FGL2
MMP19
CX3CR1
IL6
CSF3
IL1A
CCL20
AQP9
IL6
VSIR
CCL2
SOD2
FGL2
CCL7
CX3CR1
CSF3
F
MSigDB Hallmark
Monocytes
CD15⁺ myeloid
TNF-alpha Signaling via NF-kB
Inflammatory Response
Interferon Gamma Response
Interferon Alpha Response
IL-2/STAT5 Signaling
Apoptosis
Allograft Rejection
Complement
IL-6/JAK/STAT3 Signaling
KRAS Signaling Up
Epithelial Mesenchymal Transition
mTORC1 Signaling
Unfolded Protein Response
R848
G
MSigDB Hallmark
Monocytes
CD15⁺ myeloid
TNF-alpha Signaling via NF-kB
Inflammatory Response
Interferon Gamma Response
Interferon Alpha Response
IL-2/STAT5 Signaling
Apoptosis
Allograft Rejection
Complement
IL-6/JAK/STAT3 Signaling
KRAS Signaling Up
Epithelial Mesenchymal Transition
mTORC1 Signaling
UV Response Up
LPS
Gestational Age
<32+0
≥32+0
-log10(p-adj)
40
60
80
H
Monocytes
LPS response
C11orf96
TRMT10C
CSF1
PHLDA1
CHST2
THBD
FTH1
TDRD7
TRIM25
EAF1
CHMP5
SAMD9
CHSY1
ADAR
HELZ2
TUBA1C
OAS1
PLA2G4A
CNP
NCOA7
log2FC of <32+0
log2FC of ≥32+0
Significance
Higher in <32+0
Higher in ≥32+0
I
CD15⁺ myeloid cells
LPS response
TMEM158
TNFSF14
IFIT3
DNTTIP2
MAPK6
HIF1A-AS3
DDIT4
ISG15
ADAM8
DDI2
MX1
FTH1
DTX3L
IFIT1
TNRC6C
OAS2
RTCB
PARP9
OAS1
log2FC of <32+0
log2FC of ≥32+0

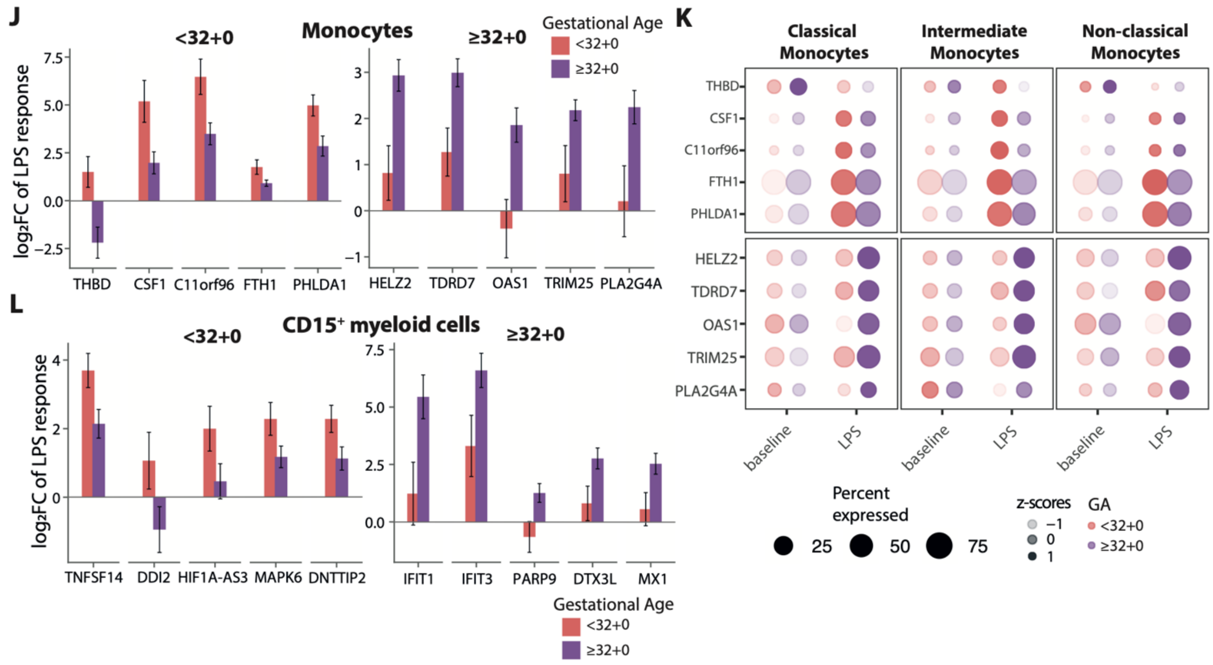


**Figure S4.**
(A) Dot plot of normalized *TLR4*, *TLR7*, and *TLR8* expression across cell types, stratified by age group. (B-E) Volcano plots of gene expression in neonatal $CD15^+$ myeloid cells and monocytes (B: R848-stimulated vs. baseline monocytes; C: R848-stimulated vs. baseline $CD15^+$ myeloid cells; D: LPS-stimulated vs. baseline monocytes; E: LPS-stimulated vs. baseline $CD15^+$ myeloid cells), stratified by age group. Thresholds are set at $|\log_2 FC| > 1$ and adjusted $p < 0.05$. (F-G) Dot plot of MSigDB enrichment (adj. $p < 0.05$) in monocytes and $CD15^+$ myeloid cells for neonates ≥32+0 and <32+0 weeks, based on genes upregulated in R848- (F) or LPS-stimulated (G) cells ($|\log_2 fC| > 1$ and adj. $p < 0.05$), ranked by adjusted p value. (H-I) Scatter plot of monocyte (H) or $CD15^+$ myeloid cell (I) $\log_2$ fold changes (LPS vs. baseline) for each gene in <32+0-wk (y-axis) and ≥32+0-wk (x-axis) neonates. Genes showing significant interaction effect (adj. $p < 0.1$) between gestational age and condition are colored. Gene labels indicate the top genes among these interaction-significant genes, ranked by adj. p. (J) Bar plots of $\log_2$ fold changes of the top five genes per age group with significant GA-dependent responses to LPS stimulation in monocytes. Error bars indicate standard errors of the DESeq2 coefficients (lfcSE) for <32+0-wk and ≥32+0-wk neonates following LPS stimulation. (K) Dot plot displaying the same genes shown in panel J as expressed by monocyte subtype. (L) Bar plots of $\log_2$ fold changes of the top five genes per age group with significant GA-dependent responses to LPS stimulation in $CD15^+$ myeloid cells, analogous to (J) for monocytes.

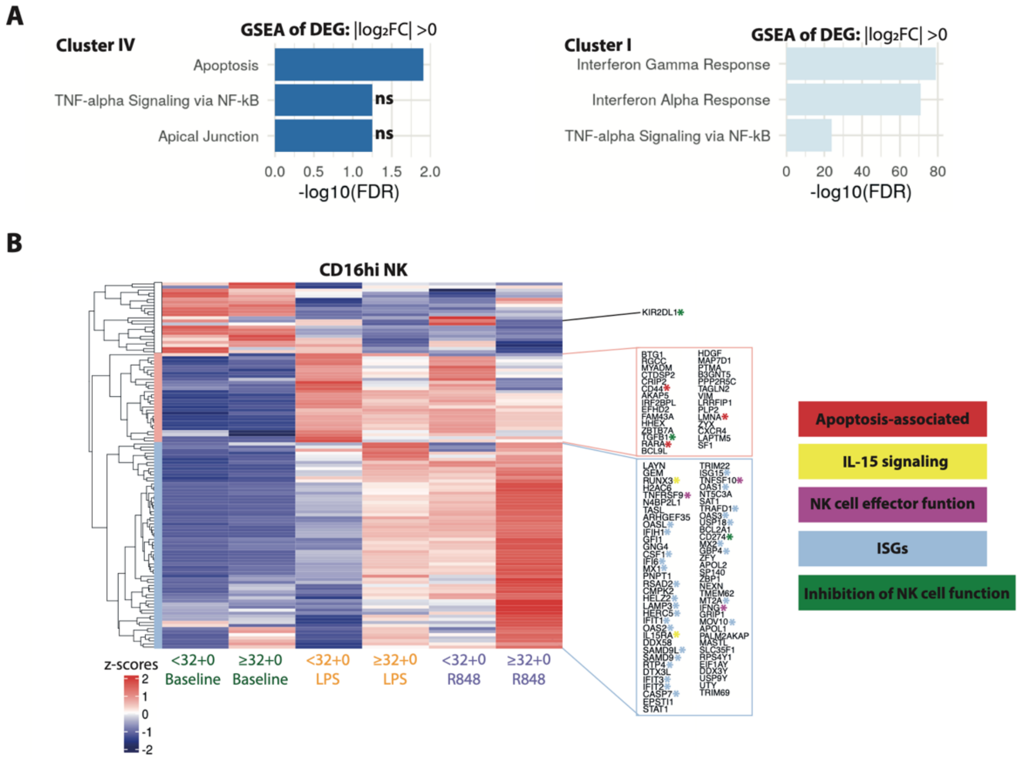


**Figure S5.**
(A) Bar plots showing the top enriched pathways based on significantly upregulated genes comparing CD16hi NK Cluster I vs. IV, ranked by adjusted p value. (B) Hierarchically clustered heatmap of top DEGs from Cluster I vs. Cluster IV (filtered based on average $\log_2$ fold change, adj. $p < 0.05$) across all CD16hi NK cells, stratified by condition and age.

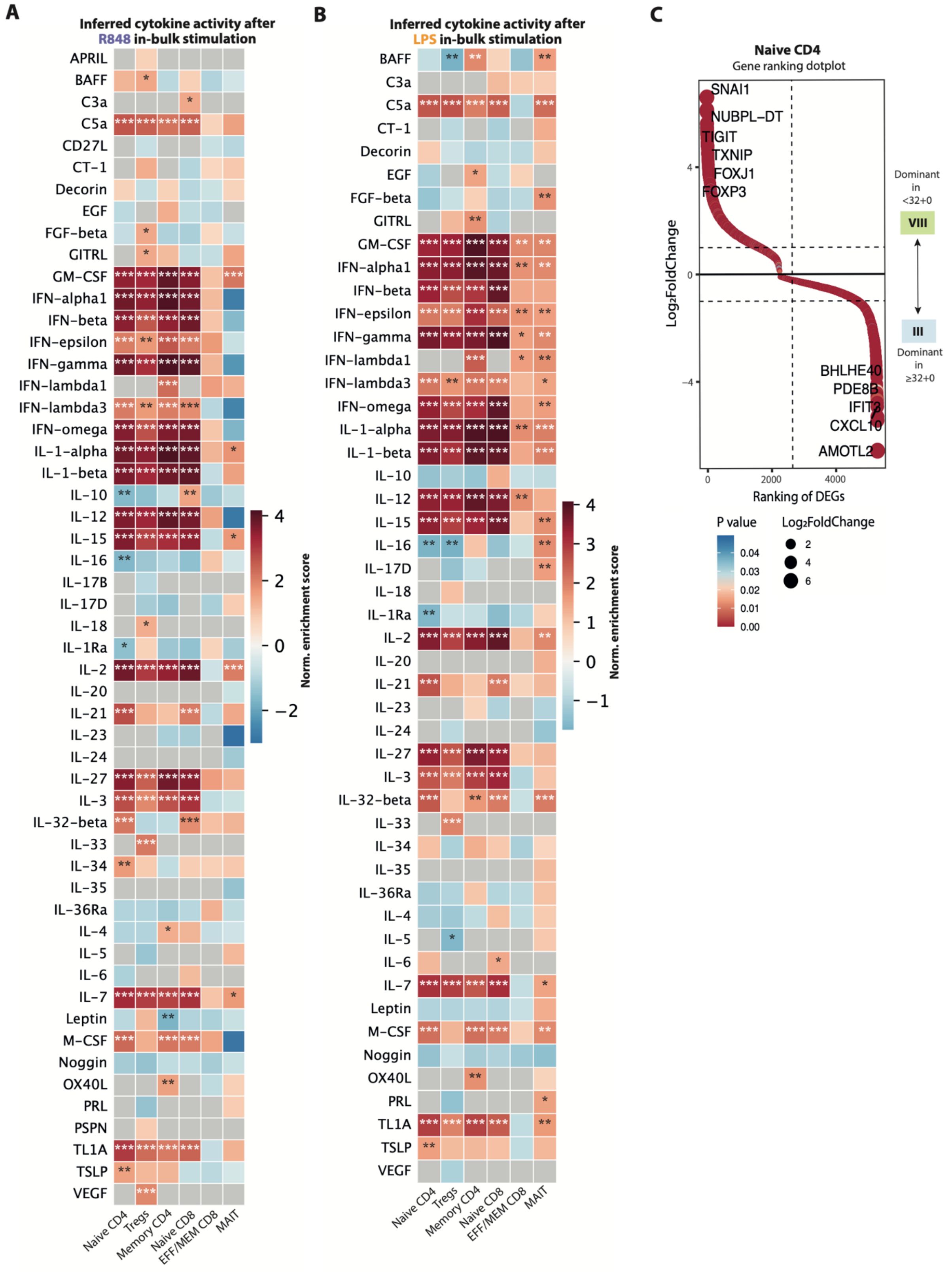


**Figure S6.**
(A-B) Heatmap showing results from huCIRA-based gene set enrichment analysis of cytokine-associated gene signatures in T cell subsets comparing ≥32+0-wk and <32+0-wk neonatal samples after R848 (A) or LPS (B) stimulation.[49] Red: enrichment in ≥32+0-wk, blue: enrichment in <32+0-wk samples. Asterisks denote statistically significant enrichment. (C) Gene ranking plots of differentially expressed genes between Cluster VIII and Cluster III naïve CD4 T cells (adj. $p < 0.05$), ranked by $\log_2$ fold change.

### Supplementary Tables

| Group | Median GA (range), weeks | Median cGA (range), weeks | Female, n (%) | Median postnatal age (range), days | Section, n (%) | Median birth weight (range), grams | Multiple gestation, n (%) |
|---|---|---|---|---|---|---|---|
| **CBMC <32+0** | 26 (24-26) | 26 (24-6) | 3 (100) | 1 | 3 (100) | 790 (690-825) | 3 (100) |
| **CBMC ≥32+0** | 38 (33-42) | 38 (33-42) | 7 (70) | 1 | 8 (80) | | 2 (20) |
| **PBMC <32+0** | 28 (26-31) | 29 (26-31) | 3 (75) | 3 (3-4) | 3 (75) | 1238 (800-1940) | 2 (50) |
| **PBMC ≥32+0** | 39 (34-41) | 39 (34-41) | 5 (63) | 3 (2-3) | 5 (63) | 3345 (1550-4100) | 0 (0) |

**Table S1. General demographics by sample group.** CBMC - cord blood mononuclear cells, PBMC - peripheral blood mononuclear cells, GA - gestational age, cGA - corrected gestational age. CBMC <32+0: n = 3, CBMC ≥32+0: n = 10, PBMC <32+0: n = 4, PBMC ≥32+0: n = 8.

| Donor-ID | GA Category | GA at Birth | cGA | Sex | Postnatal Age (days) | Mode of Delivery | Birth weight (g) | Multiple gestation |
|---|---|---|---|---|---|---|---|---|
| **CBMC1** | extreme | 26(2/7) | 26(2/7) | female | 1 | section | 790 | yes |
| **CBMC2** | extreme | 26(2/7) | 26(2/7) | female | 1 | section | 825 | yes |
| **CBMC3** | extreme | 24(3/7) | 24(3/7) | female | 1 | section | 690 | yes |
| **CBMC4** | late | 33(1/7) | 33(1/7) | female | 1 | section | | no |
| **CBMC5** | late | 35(2/7) | 35(2/7) | female | 1 | vaginal | | yes |
| **CBMC6** | late | 35(2/7) | 35(2/7) | female | 1 | vaginal | | yes |
| **CBMC7** | term | 42(0/7) | 42(0/7) | male | 1 | section | | no |
| **CBMC8** | term | 39(0/7) | 39(0/7) | male | 1 | section | | no |
| **CBMC9** | term | 38(4/7) | 38(4/7) | female | 1 | section | | no |
| **CBMC10** | term | 39(0/7) | 39(0/7) | female | 1 | section | | no |
| **CBMC11** | term | 39(0/7) | 39(0/7) | female | 1 | section | | no |
| **CBMC12** | term | 37(0/7) | 37(0/7) | male | 1 | section | | no |
| **CBMC13** | term | 39(0/7) | 39(0/7) | female | 1 | section | | no |
| **PBMC1** | extreme | 26(6/7) | 27(1/7) | female | 3 | section | 800 | no |
| **PBMC2** | extreme | 26(2/7) | 26(5/7) | female | 4 | section | 825 | yes |
| **PBMC3** | very | 31(4/7) | 31(6/7) | male | 3 | vaginal | 1940 | no |
| **PBMC4** | very | 31(0/7) | 31(2/7) | female | 3 | section | 1650 | yes |
| **PBMC5** | late | 35(3/7) | 35(5/7) | female | 3 | section | 3130 | no |
| **PBMC6** | late | 34(2/7) | 34(3/7) | male | 2 | section | 1550 | no |
| **PBMC7** | term | 39(4/7) | 39(6/7) | female | 3 | section | 3025 | no |
| **PBMC8** | term | 39(5/7) | 40(0/7) | male | 3 | section | 3560 | no |
| **PBMC9** | term | 40(5/7) | 41(0/7) | male | 3 | section | 4100 | no |
| **PBMC10** | term | 38(2/7) | 38(4/7) | female | 3 | vaginal | 2810 | no |
| **PBMC11** | term | 41(3/7) | 41(5/7) | female | 3 | vaginal | 3710 | no |
| **PBMC12** | term | 40(2/7) | 40(4/7) | female | 3 | vaginal | 3725 | no |

**Table S2. General demographics by infant.** CBMC - cord blood mononuclear cells, PBMC - peripheral blood mononuclear cells, GA - gestational age, cGA - corrected gestational age.

| Group | PROM | Hypertension | Preeclampsia | IUGR | GD | ACS | Maternal Antibiotics |
|---|---|---|---|---|---|---|---|
| **CBMC <32+0** | 1 (33) | 0 (0) | 0 (0) | 0 (0) | 0 (0) | 3 (100) | 3 (100) |
| **CBMC ≥32+0** | | | | | | | |
| **PBMC <32+0** | 0 (0) | 0 (0) | 1 (25) | 1 (25) | 0 (0) | 2 (50) | 1 (25) |
| **PBMC ≥32+0** | 1 (13) | 0 (0) | 1 (13) | 1 (13) | 2 (25) | 0 (0) | 0 (0) |

**Table S3. Prenatal demographics by sample group, n (%).** CBMC - cord blood mononuclear cells, PBMC - peripheral blood mononuclear cells, PROM - premature rupture of membranes, IUGR - intrauterine growth restriction, GD - gestational diabetes mellitus, ACS - antenatal corticosteroid therapy, completed <7 d prior to delivery, CBMC <32+0: n = 3, CBMC ≥32+0: n = 10, PBMC <32+0: n = 4, PBMC ≥32+0: n = 8.

| Donor-ID | GA Category | PROM | Hypertension | Preeclampsia | IUGR | GD | ACS | Maternal Antibiotics |
|---|---|---|---|---|---|---|---|---|
| **CBMC1** | extreme | yes | no | no | no | no | yes | yes |
| **CBMC2** | extreme | no | no | no | no | no | yes | yes |
| **CBMC3** | extreme | no | no | no | no | no | yes | yes |
| **CBMC4** | late | | | | | | | |
| **CBMC5** | late | | | | | | | |
| **CBMC6** | late | | | | | | | |
| **CBMC7** | term | | | | | | | |
| **CBMC8** | term | | | | | | | |
| **CBMC9** | term | | | | | | | |
| **CBMC10** | term | | | | | | | |
| **CBMC11** | term | | | | | | | |
| **CBMC12** | term | | | | | | | |
| **CBMC13** | term | | | | | | | |
| **PBMC1** | extreme | no | no | yes | yes | no | yes | no* |
| **PBMC2** | extreme | no | no | no | no | no | yes | yes |
| **PBMC3** | very | no | no | no | no | no | no | no |
| **PBMC4** | very | no | no | no | no | no | no | no* |
| **PBMC5** | late | yes | no | no | no | yes | no | no* |
| **PBMC6** | late | no | no | yes | yes | yes | no | no* |
| **PBMC7** | term | NA | no | no | no | no | no | no* |
| **PBMC8** | term | no | no | no | no | no | no | no* |
| **PBMC9** | term | no | no | no | no | no | no | no* |
| **PBMC10** | term | no | no | no | no | no | no | no |
| **PBMC11** | term | no | no | no | no | no | no | no |
| **PBMC12** | term | no | no | no | no | no | no | no |

**Table S4. Prenatal demographics by infant.** CBMC - cord blood mononuclear cells, PBMC - peripheral blood mononuclear cells, PROM - premature rupture of membranes, IUGR - intrauterine growth restriction, GD - gestational diabetes mellitus, ACS - antenatal corticosteroid therapy, completed <7 d prior to delivery, CBMC <32+0: n = 3, CBMC ≥32+0: n = 10, PBMC <32+0: n = 4, PBMC ≥32+0: n = 8. *perioperative prophylaxis for cesarean section.

| Group | SGA | AIS | EOS | Antibiotics | PDA | BPD | FIP | NEC | IVH |
|---|---|---|---|---|---|---|---|---|---|
| **CBMC <32+0** | 0 (0) | 0 (0) | 0 (0) | 3 (100) | 2 (67) | 2 (67) | 0 (0) | 0 (0) | 1 (33) |
| **CBMC ≥32+0** | | 1 (10) | | | | | | | |
| **PBMC <32+0** | 0 (0) | 0 (0) | 0 (0) | 4 (100) | 2 (50) | 2 (50) | 0 (0) | 0 (0) | 0 (0) |
| **PBMC ≥32+0** | 1 (13) | 0 (0) | 0 (0) | 0 (0) | 0 (0) | 0 (0) | 0 (0) | 0 (0) | 0 (0) |

**Table S5. Neonatal complications by sample group, n (%).** CBMC - cord blood mononuclear cells, PBMC - peripheral blood mononuclear cells, SGA - small for gestational age, AIS - amniotic infection syndrome, EOS - early-onset sepsis, PDA - patent ductus arteriosus, BPD - Bronchopulmonary Dysplasia, FIP - focal intestinal perforation, NEC - necrotizing enterocolitis, IVH - intraventricular hemorrhage, CBMC <32+0: n = 3, CBMC ≥32+0: n = 10, PBMC <32+0: n = 4, PBMC ≥32+0: n = 8.

| Donor-ID | GA Category | SGA | AIS | EOS | Antibiotics | PDA | BPD | FIP | NEC | IVH |
|---|---|---|---|---|---|---|---|---|---|---|
| **CBMC1** | extreme | no | no | Pneu-monia | yes | no | no | no | no | Gr. I |
| **CBMC2** | extreme | no | no | no | yes | yes | mild | no | no | no |
| **CBMC3** | extreme | no | no | no | yes | yes | mod | no | no | no |
| **CBMC4** | late | | yes* | | | | | | | |
| **CBMC5** | late | | no | | | | | | | |
| **CBMC6** | late | | no | | | | | | | |
| **CBMC7** | term | | no | | | | | | | |
| **CBMC8** | term | | no | | | | | | | |
| **CBMC9** | term | | no | | | | | | | |
| **CBMC10** | term | | no | | | | | | | |
| **CBMC11** | term | | no | | | | | | | |
| **CBMC12** | term | | no | | | | | | | |
| **CBMC13** | term | | no | | | | | | | |
| **PBMC1** | extreme | no | no | no | yes | yes | mild | no | no | no |
| **PBMC2** | extreme | no | no | no | yes | yes | mild | no | no | no |
| **PBMC3** | very | no | no | no | yes | no | no | no | no | no |
| **PBMC4** | very | no | no | no | yes | no | no | no | no | no |
| **PBMC5** | late | no | no | no | no | no | no | no | no | no |
| **PBMC6** | late | yes | no | no | no | no | no | no | no | no |
| **PBMC7** | term | no | no | no | no | no | no | no | no | no |
| **PBMC8** | term | no | no | no | no | no | no | no | no | no |
| **PBMC9** | term | no | no | no | no | no | no | no | no | no |
| **PBMC10** | term | no | no | no | no | no | no | no | no | no |
| **PBMC11** | term | no | no | no | no | no | no | no | no | no |
| **PBMC12** | term | no | no | no | no | no | no | no | no | no |

**Table S6. Neonatal complications by infant.** CBMC - cord blood mononuclear cells, PBMC - peripheral blood mononuclear cells, SGA - small for gestational age, AIS - amniotic infection syndrome, EOS - early-onset sepsis, PDA - patent ductus arteriosus, BPD - Bronchopulmonary Dysplasia, FIP - focal intestinal perforation, NEC - necrotizing enterocolitis, IVH - intraventricular hemorrhage. Gr. - Grade, mod - moderate. *Clinically diagnosed; no placental histology available.

| Donor-ID | GA Cat. | WBC, G/l | Hb, g/dl | HCT, l/l | PLT, G/l | Neut, G/l | Lymph, G/l | Mono, G/l | Eosino, G/l | Baso, G/l | CrP, mg/dl | IL-6, pg/ml |
|---|---|---|---|---|---|---|---|---|---|---|---|---|
| **CBMC1** | extreme | 20.1 | 16.8 | 0.480 | 275 | 13.8 | 3.07 | 2.06 | 0.11 | 0.11 | <0.1 | 351 |
| **CBMC2** | extreme | 7.86 | 14.5 | 0.421 | 280 | 1.62 | 5.05 | 0.64 | 0.45 | 0.04 | <0.1 | 128 |
| **CBMC3** | extreme | 4.56 | 13.6 | 0.378 | 182 | 0.27 | NA | NA | NA | NA | <0.1 | 180 |
| **CBMC4** | late | | | | | | | | | | | |
| **CBMC5** | late | | | | | | | | | | | |
| **CBMC6** | late | | | | | | | | | | | |
| **CBMC7** | term | | | | | | | | | | | |
| **CBMC8** | term | | | | | | | | | | | |
| **CBMC9** | term | | | | | | | | | | | |
| **CBMC10** | term | | | | | | | | | | | |
| **CBMC11** | term | | | | | | | | | | | |
| **CBMC12** | term | | | | | | | | | | | |
| **CBMC13** | term | | | | | | | | | | | |
| **PBMC1** | extreme | 6.69 | 15.5 | 0.467 | 225 | 1.72 | 3.31 | 1.40 | 0.15 | 0.02 | 0.1 | 24.5 |
| **PBMC2** | extreme | 9.93 | 12.0 | 0.342 | 220 | 4.48 | 2.87 | 0.82 | 1.65 | 0.07 | 0.7 | NA |
| **PBMC3** | very | NA | NA | NA | NA | NA | NA | NA | NA | NA | NA | NA |
| **PBMC4** | very | 14.2 | 16.8 | 0.492 | 370 | NA | NA | NA | NA | NA | 0.1 | NA |
| **PBMC5** | late | 7.85 | 13.8 | 0.395 | 221 | 2.88 | 3.86 | 0.77 | 0.29 | 0.02 | <0.1 | 21.8 |
| **PBMC6** | late | 8.81 | 21.2 | 0.613 | 141 | NA | NA | NA | NA | NA | <0.1 | <22.5 |
| **PBMC7** | term | NA | NA | NA | NA | NA | NA | NA | NA | NA | NA | NA |
| **PBMC8** | term | NA | NA | NA | NA | NA | NA | NA | NA | NA | NA | NA |
| **PBMC9** | term | NA | NA | NA | NA | NA | NA | NA | NA | NA | NA | NA |
| **PBMC10** | term | NA | NA | NA | NA | NA | NA | NA | NA | NA | NA | NA |
| **PBMC11** | term | NA | NA | NA | NA | NA | NA | NA | NA | NA | NA | NA |
| **PBMC12** | term | NA | NA | NA | NA | NA | NA | NA | NA | NA | NA | NA |

**Table S7. Complete blood count data (absolute counts) and infectious parameters within 24 hours of blood draw.** CBMC - cord blood mononuclear cells, PBMC - peripheral blood mononuclear cells, GA - gestational age, DOL - day of life, WBC - white blood cell count, Hb - Hemoglobin, HCT - hematocrit, PLT - platelets, Neut - Neutrophils, Lymph - Lymphocytes, Mono - Monocytes, Eosino - Eosinophils, Baso - Basophils.

| Specificity | Clone | Oligo ID | Specificity | Clone | Oligo ID | Specificity | Clone | Oligo ID |
|---|---|---|---|---|---|---|---|---|
| CD3 | UCHT1 | AHS0231 | CD45RA | HI100 | AHS0009 | CD196 (CCR6) | 11A9 | AHS0034 |
| CD4 | SK3 | AHS0032 | CD56 | NCAM16 | AHS0019 | CD197 (CCR7) | 2-L1-A | AHS0273 |
| CD8 | SK1 | AHS0228 | CD62L | DREG-56 | AHS0049 | CD272 | J168-540 | AHS0052 |
| CD11c | B-Ly6 | AHS0056 | CD127 | HIL-7R-M21 | AHS0028 | CD278 | DX29 | AHS0012 |
| CD14 | MPHIP9 | AHS0037 | CD134 | ACT35 | AHS0013 | CD279 | EH12.1 | AHS0014 |
| CD16 | 3G8 | AHS0053 | CD137 | 4B4-1 | AHS0003 | CD357 (GITR) | V27-580 | AHS0104 |
| CD19 | SJ25C1 | AHS0030 | CD161 | HP-3G10 | AHS0205 | CD366 (TIM-3) | 7D3 | AHS0016 |
| CD25 | 2A3 | AHS0026 | CD183 (CXCR3) | 1C6/CXCR3 | AHS0031 | HLA-DR | G46-6 | AHS0035 |
| CD27 | M-T271 | AHS0025 | CD185 (CXCR5) | RF8B2 | AHS0039 | IgD | IA6-2 | AHS0058 |

| | | | | | | | | |
|---|---|---|---|---|---|---|---|---|
| CD28 | L293 | AHS0138 | CD186 (CXCR6) | 13B 1E5 | AHS0148 | IgM | G20-127 | AHS0198 |

**Table S8. BD AbSeq Immune Discovery Panel.** List of BD AbSeq IDP Specificities

| Specificity | Clone | Oligo ID |
|---|---|---|
| CRTH2 | BM16 | AHS0106 |
| CD117 | 104D2 | AHS0165 |
| γδ TCR | B1 | AHS0015 |
| CD15 | W6D3 | AHS0196 |
| CD141 | 1A4 | AHS0083 |
| CD11b | ICRF44 | AHS0184 |

**Table S9. Custom Spike-In BD AbSeq specificities added to the BD AbSeq Immune Discovery Panel**

| Group | Median GA (range), weeks | Median cGA (range), weeks | Female, n (%) | Median postnatal age (range), days | Section, n (%) | Median birth weight (range), grams | Multiple gestation, n (%) |
|---|---|---|---|---|---|---|---|
| **PBMC <32+0** | 28 (23-31) | 28 (23-31) | 8 (38) | 3 (1-7) | 16 (76) | 970 (620-1820) | 6 (29) |
| **PBMC ≥32+0** | 37 (26-41) | 38 (32-42) | 20 (63) | 3 (3-37) | 16 (52) | 2605 (800-4175) | 4 (13) |

**Table S10. General demographics by sample group (flow cytometry validation cohort).** CBMC - cord blood mononuclear cells, PBMC - peripheral blood mononuclear cells, GA - gestational age, cGA - corrected gestational age. PBMC <32+0: n = 21, PBMC ≥32+0: n = 32.

| Donor-ID | GA Category | GA at Birth | cGA | Sex | Postnatal Age (days) | Mode of Delivery | Birth weight (g) | Multiple gestation |
|---|---|---|---|---|---|---|---|---|
| **PBMC13** | extreme | 26(2/7) | 26(4/7) | male | 3 | section | 970 | no |
| **PBMC14** | extreme | 26(1/7) | 26(2/7) | male | 2 | vaginal | 970 | no |
| **PBMC15** | extreme | 23(3/7) | 23(5/7) | male | 3 | vaginal | 675 | yes |
| **PBMC16** | extreme | 23(3/7) | 23(5/7) | male | 3 | vaginal | 620 | yes |
| **PBMC17** | extreme | 24(3/7) | 24(5/7) | female | 3 | section | 650 | yes |
| **PBMC18** | extreme | 24(3/7) | 24(5/7) | female | 3 | section | 690 | yes |
| **PBMC19** | extreme | 26(6/7) | 27(3/7) | male | 5 | section | 740 | no |
| **PBMC20** | extreme | 25(3/7) | 26(0/7) | male | 5 | section | 745 | no |
| **PBMC21** | very | 29(5/7) | 30(0/7) | female | 3 | section | 1060 | no |
| **PBMC22** | very | 29(1/7) | 29(4/7) | female | 4 | vaginal | 1320 | no |
| **PBMC23** | very | 31(2/7) | 31(4/7) | male | 3 | vaginal | 1820 | no |
| **PBMC24** | very | 27(6/7) | 28(2/7) | male | 4 | section | 850 | yes |
| **PBMC25** | very | 31(0/7) | 31(2/7) | male | 3 | section | 1795 | yes |
| **PBMC26** | very | 31(0/7) | 31(2/7) | male | 3 | section | 1580 | no |
| **PBMC27** | very | 28(4/7) | 28(5/7) | male | 2 | section | 1240 | no |
| **PBMC28** | very | 29(1/7) | 29(3/7) | male | 3 | section | 1340 | no |
| **PBMC29** | very | 29(0/7) | 29(6/7) | female | 7 | section | 940 | no |
| **PBMC30** | very | 28(3/7) | 29(0/7) | female | 5 | section | 1355 | no |
| **PBMC31** | very | 28(2/7) | 29(1/7) | female | 7 | section | 805 | no |
| **PBMC32** | very | 29(2/7) | 29(2/7) | male | 1 | section | 1050 | no |
| **PBMC33** | very | 28(0/7) | 28(3/7) | female | 4 | section | 1000 | no |

| | | | | | | | | |
|---|---|---|---|---|---|---|---|---|
| **PBMC34** | late | 33(3/7) | 33(5/7) | female | 3 | section | 1730 | no |
| **PBMC35** | late | 32(1/7) | 32(5/7) | male | 5 | section | 1030 | no |
| **PBMC36** | late | 32(6/7) | 33(1/7) | female | 3 | section | 1860 | no |
| **PBMC37** | late | 32(1/7) | 32(3/7) | female | 3 | vaginal | 1800 | no |
| **PBMC38** | late | 34(1/7) | 34(3/7) | male | 3 | section | 2200 | yes |
| **PBMC39** | late | 34(1/7) | 34(3/7) | male | 3 | section | 1970 | yes |
| **PBMC40** | late | 33(3/7) | 34(4/7) | male | 9 | section | 1300 | no |
| **PBMC41** | late | 31(1/7) | 32(0/7) | female | 7 | section | 1740 | no |
| **PBMC42** | late | 34(5/7) | 35(0/7) | female | 3 | section | 2280 | no |
| **PBMC43** | late | 33(2/7) | 33(4/7) | female | 3 | vaginal | 2165 | no |
| **PBMC44** | late | 29(3/7) | 32(3/7) | female | 22 | section | 890 | no |
| **PBMC45** | late | 26(6/7) | 32(0/7) | female | 37 | section | 800 | no |
| **PBMC46** | late | 33(1/7) | 33(5/7) | female | 5 | section | 1300 | no |
| **PBMC47** | late | 33(2/7) | 33(4/7) | female | 3 | section | 1215 | no |
| **PBMC48** | term | 39(6/7) | 40(1/7) | male | 3 | vaginal | 3475 | no |
| **PBMC49** | term | 39(6/7) | 40(1/7) | female | 3 | vaginal | 3970 | no |
| **PBMC50** | term | 41(1/7) | 41(3/7) | female | 3 | vaginal | 3100 | no |
| **PBMC51** | term | 39(5/7) | 40(0/7) | female | 3 | section | 3650 | no |
| **PBMC52** | term | 41(3/7) | 41(5/7) | female | 3 | vaginal | 3430 | no |
| **PBMC53** | term | 39(5/7) | 40(0/7) | male | 3 | vaginal | 3170 | no |
| **PBMC54** | term | 39(3/7) | 39(5/7) | male | 3 | vaginal | 3465 | no |
| **PBMC55** | term | 39(3/7) | 39(5/7) | female | 3 | section | 3355 | no |
| **PBMC56** | term | 39(3/7) | 39(5/7) | male | 3 | section | 3195 | no |
| **PBMC57** | term | 40(2/7) | 40(4/7) | male | 3 | vaginal | 3355 | no |
| **PBMC58** | term | 41(5/7) | 42(0/7) | male | 3 | vaginal | 4025 | no |
| **PBMC59** | term | 41(5/7) | 42(0/7) | male | 3 | vaginal | 4175 | no |
| **PBMC60** | term | 37(3/7) | 37(5/7) | female | 3 | vaginal | 2605 | yes |
| **PBMC61** | term | 37(3/7) | 37(5/7) | female | 3 | vaginal | 1945 | yes |
| **PBMC62** | term | 39(3/7) | 39(5/7) | female | 3 | vaginal | 2825 | no |
| **PBMC63** | term | 39(3/7) | 39(5/7) | male | 3 | | | |
| **PBMC64** | term | 36(6/7) | 37(1/7) | female | 3 | vaginal | 2880 | no |
| **PBMC65** | term | 38(4/7) | 38(6/7) | female | 3 | vaginal | 3030 | no |

**Table S11. General demographics by infant at the time of blood draw (flow cytometry validation cohort).** CBMC - cord blood mononuclear cells, PBMC - peripheral blood mononuclear cells, GA - gestational age, cGA - corrected gestational age.

| score | reference | compared_to | p_value | cliff_delta | p_adj |
|---|---|---|---|---|---|
| **Th1_score** | III | II | 1.76944871259175e-49 | 0.168664841927748 | 6.63543267221906e-49 |
| **Th1_score** | III | VIII | 5.53784958567565e-09 | 0.0816092604840286 | 9.22974930945941e-09 |
| **Th1_score** | III | I | 1.66179669377524e-35 | 0.137446174111343 | 4.98539008132573e-35 |
| **Th2_score** | VIII | II | 0.0136315863578721 | 0.0305837366442792 | 0.0185885268516438 |
| **Th2_score** | VIII | III | 0.000206353559296129 | 0.0504482106195952 | 0.000309530338944193 |
| **Th2_score** | VIII | I | 0.999999999201071 | -0.081901524579387 | 1 |

| | | | | | |
|---|---|---|---|---|---|
| **Th2_score** | I | II | 2.6058561822582e-29 | 0.117836543326475 | 5.58397753341043e-29 |
| **Th2_score** | I | III | 3.87996413107777e-35 | 0.136688654465549 | 9.69991032769442e-35 |
| **Th2_score** | I | VIII | 7.98973926747828e-10 | 0.081901524579387 | 1.49807611265218e-09 |
| **Treg_score** | I | II | 3.55214900672245e-219 | 0.332896536127007 | 2.66411175504183e-218 |
| **Treg_score** | I | III | 0 | 0.439619815417003 | 0 |
| **Treg_score** | I | VIII | 5.80742740332521e-199 | 0.408152633078476 | 2.9037137016626e-198 |
| **Treg_score** | VIII | II | 0.999999999994941 | -0.0942668277266407 | 1 |
| **Treg_score** | VIII | III | 0.344095291590969 | 0.00573237523147288 | 0.430119114488711 |
| **Treg_score** | VIII | I | 1 | -0.408152633078476 | 1 |

**Table S12. CD4+ T cell gene scores.** Single-cell module scores for Th1, Th2, and Treg gene sets as computed using Seurat's AddModuleScore on normalized data (calculated at the single-cell level; pairwise Wilcoxon rank-sum tests). Gene sets were derived from published T helper signatures[9] and the MSigDB C7 immunologic signature collection.[70]

| Conjugate | Specificity | Clone | Manufacturer |
|---|---|---|---|
| Pacific Blue | CD3 | SK7 | BioLegend |
| FITC | CD4 | RPA-T4 | BioLegend |
| APC/Fire 750 | CD8 | HIT8a | BioLegend |
| PE-CF594 | CD56 | NCAM16.2 | BD Biosciences |
| R718 | CD19 | HIB19 | BD Biosciences |
| FVS | Live/Dead | | BD Biosciences |

**Table S13. Flow cytometry antibodies (Panel 1)**

| Conjugate | Specificity | Clone | Manufacturer |
|---|---|---|---|
| BUV496 | CD3 | UCHT1 | BD Biosciences |
| PE-Cy5 | CD8 | RPA-T8 | BD Biosciences |
| R718 | CD19 | HIB19 | BD Biosciences |
| BB700 | CD56 | NCAM16.2 | BD Biosciences |
| BV785 | CD14 | M5E2 | BioLegend |
| BV605 | CD16 | 3G8 | BioLegend |
| BB515 | HLA-DR | G46-6 | BD Biosciences |
| APC | CD11c | B-ly6 | BD Biosciences |
| BV421 | CD123 | 6H6 | BD Biosciences |
| BV711 | CD5 | UCHT2 | BD Biosciences |
| PE/Dazzle 594 | CD1c | L161 | BioLegend |
| BUV737 | CD141 | 1A4 | BD Biosciences |
| PE | Axl | 108724 | R&D Systems |
| BV480 | CD45RA | HI100 | BD Biosciences |
| PE-Cy7 | CD303 | 201A | BioLegend |
| FVS | Live/Dead | | BD Biosciences |

**Table S14. Flow cytometry antibodies (Panel 2)**

| Conjugate | Target | Clone | Manufacturer |
|---|---|---|---|
| BUV496 | CD3 | UCHT1 | BD Biosciences |
| APC/Fire 750 | CD8 | HIT8a | BioLegend |
| FITC | CD4 | RPA-T4 | BioLegend |
| APC | CD62L | 145/15 | Milteny |
| BV480 | CD45RA | HI100 | BD Biosciences |
| PE-CF594 | CD56 | NCAM16.2 | BD Biosciences |
| BV605 | CD16 | 3G8 | BioLegend |
| BV785 | CD14 | M5E2 | BioLegend |
| R718 | CD19 | HIB19 | BD Biosciences |
| PE | HLA-DR | L243 | BioLegend |
| BV421 | TCRyd | 11F2 | BD Biosciences |
| RB705 | CD11b | ICRF44 | BD Biosciences |
| BV711 | CD10 | HI10a | BD Biosciences |
| PE-Cy7 | CD15 | HI98 | BD Biosciences |
| 7-AAD | Live/dead | | BD Biosciences |

**Table S15. Flow cytometry antibodies (Panel 3)**

**Table S16.** Differentially expressed genes between neonatal and adult immune cells by cell type as assessed using DESeq2 (adj. $p < 0.05$). For each cell type, the table lists gene symbol, $\log_2$ FC (adult vs. neonatal), and adjusted p value.

**Table S17.** Differentially expressed genes between birth and DOL3 neonatal immune cells by cell type as assessed using DESeq2 (adj. $p < 0.05$). For each cell type, the table lists gene symbol, $\log_2$ FC (DOL3 vs. birth), and adjusted p value.

**Table S18.** Differentially expressed genes between LPS and baseline neonatal immune cells, as well as R848 and baseline immune cells by cell type as assessed using DESeq2 (adj. $p < 0.05$, $|\log_2 FC| > 1$). For each cell type, the table lists gene symbol, $\log_2$ FC (stim vs. baseline), and adjusted p value. To test for differential responses between the two GA groups, an interaction analysis was performed only for monocytes and $CD15^+$ myeloid cells, modeling gene expression as a function of GA group, stimulation, and their interaction (GA group × stimulation). For these two cell types, the adjusted p value for the interaction term is additionally reported.
